# Inferring cell trajectories of spatial transcriptomics via optimal transport analysis

**DOI:** 10.1101/2023.09.04.556175

**Authors:** Xunan Shen, Ke Huang, Lulu Zuo, Zhongfei Ye, Zeyu Li, Qichao Yu, Xuanxuan Zou, Xiaoyu Wei, Ping Xu, Xin Jin, Xun Xu, Liang Wu, Hongmei Zhu, Pengfei Qin

**Author notes:** Correspondence (P.Q.), (H.Z.), (L.W.), (X.X). These authors contributed equally to this work (Xunan Shen, Ke Huang, Lulu Zuo, Zhongfei Ye).

## Abstract

The integration of cell transcriptomics and spatial coordinates to organize differentiation trajectories remains a challenge. Here we introduce spaTrack, a trajectory inference method using optimal transport to incorporate both transcriptomics and distance of spatial transcriptomics sequencing data into transition costs. spaTrack could construct fine spatial trajectories reflecting the true differentiation topology, as well as trace cell dynamics across multiple samples with temporal intervals. To capture the dynamic drivers, spaTrack models the cell fate as a function of expression profile along temporal intervals driven by transcription factors. Applying spaTrack, we successfully disentangle spatiotemporal trajectories of axolotl telencephalon regeneration and mouse midbrain development. Furthermore, we uncover diverse malignant lineages expanding in a primary tumor. One of the lineages with upregulated extracellular matrix organization implants to the metastatic site and subsequently colonizes to a secondary tumor. Overall, spaTrack greatly facilitates trajectory inference from spatial transcriptomics, providing insights in cell differentiation of broad areas.

## Introductions

Trajectory inference (TI) provides important insights in cell differentiation and biological process. Currently, there are numerous TI methods available, but most of them are designed for single cell RNA sequencing (SC) data and challenged by complex topologies. Frequently applied TI methods for SC data, e.g., Monocle2/3 (Cao et al., 2019; Qiu et al., 2017), PAGA (Wolf et al., 2019), Slingshot (Street et al., 2018), stemID (Grün et al., 2016), Tscan (Ji and Ji, 2016), URD (Farrell et al., 2018) et al. usually construct a skeleton frame of cell differentiation averaged or optimally extracted from a SC data embedding generated by dimension reduction (DR) such as PCA, ICA, UMAP (McInnes et al., 2018), Diffusion Map (Haghverdi et al., 2015) and ForceAtlas2 (Jacomy et al., 2014). Many of the existing approaches are limited to simple linear or branched topologies and overlook morphometric space. However, the dynamics of biological systems, such as embryonic development or tumor progression, are often complex and strictly spatially organized. Cell transition is spatially heterogenous due to their location and surrounding environment. Recent advances in spatial transcriptomics sequencing (ST) technologies provide an opportunity to simultaneously reveal both transcriptomic and spatial patterns of development, which the SC is unable to capture. Trajectories generated from SC data and their methods are uncapable to uncover the spatial details of differentiation, and discrete trajectories are often compelled to be continuous in the SC manner, conflicting with the true topology.

To capture the single cell dynamics, RNA velocity has introduced alternative ways to study cellular differentiation of SC data (Bergen et al., 2020; Chen et al., 2022c; La Manno et al., 2018; Qiu et al., 2022). It describes the rate of transcriptional dynamics for an individual gene at a given time point based on the ratio of its spliced and unspliced messenger RNA (mRNA). Great efforts have been made to develop various TI methods based on RNA velocity. However, estimation of RNA velocity is often found to be less robust for indicating cellular transitions due to several fundamental limitations. There are low contents of unspliced RNAs in SC data and intronic regions could not be fully captured; Conventional RNA-velocity methods assume constant transcription rates but the cell population/states are heterogeneous which usually leads to nonsensical backward trajectories. Metabolic-labeling data has been explored in velocity estimate, which measures the synthesis and degradation of labeled RNA within a known period of time directly and overcomes some drawbacks of conventional RNA-velocity methods (Qiu et al., 2022). However, the metabolic-labeling data is not always available in studies. In addition, most RNA-velocity methods do not reconcile the physical proximity of cells into trajectory inference. Besides, estimating RNA-splicing rate involves mapping BAM file which can be computationally time-consuming and require significant computational resources. Thus, there is a strong need for an alternative approach to efficiently generate single-cell spatial trajectories.

Optimal transport (OT) is a method to find least-cost schemes of coupling distributions and provide intuitively quantifications of their distance between multiple datasets or samples represented as distributions (Villani, 2009). OT has recently been used for transcriptomic data analysis, including cell-cell communication inference (Cang and Nie, 2020; Cang et al., 2023), lineage study of hematogenesis (Schiebinger et al., 2019) and annotation of ST data (Cang and Nie, 2020; Nitzan et al., 2019). With the natural advantage of OT, it is capable to conveniently incorporate the profile of both gene expression and transition distance of cells into the cost matrix to solve the OT problem. A fully captured cell to cell transition matrix will facilitate the construction of trajectories with fine local details.

In this study, we introduced an efficient method, namely spaTrack, to construct cell trajectory at single cell resolution of spatial context, which utilizes OT frameworks and sensitively reconciles both gene expression and physical distance. When dealing with multiple samples from a time series, spaTrack can construct a dynamic map of cell migration and differentiation across all tissue sections, providing a comprehensive view of transition behavior over time. In our study, spaTrack performs reliably in various scenarios of SC and ST data. We have successfully applied spaTrack to reconstruct cell trajectories in spatial manners for various biological systems, including regeneration of injured axolotl telencephalon, development of dorsal midbrain of mouse embryo, and tumor expansion and metastasis. Our approach has significantly facilitated the study of cell kinetics of ST data in a wide range of cases.

## Results

### Inferring cell trajectories from single ST data

spaTrack utilizes optimal transport (OT) as a foundation to infer the transition probability between cells of ST data in a single sample, by incorporating both gene expression profiles and cell location information. Cells that are distant from each other in expression level and physical space will have higher transfer costs, which indicates a lower transition probability or longer time interval in the biological process. The schematics of the algorithm and workflow of spaTrack in solving the TI problem in single ST data are demonstrated in Figure 1A, which include 1) scaling both gene expression and physical distance into a cost matrix; 2) solving the OT problem by incorporating an entropy term; 3) constructing the vector field of cell velocity from transferring probability; 4) organizing spatial trajectories from the vector field of cell velocity; 5) optimizing the path between a starting cell and corresponding ending cell using the least action path method; 6) identifying pseudotime-dependent genes using a generalized additive model (GAM) to fit the dynamics of gene expression along a trajectory.

**Figure 1.**
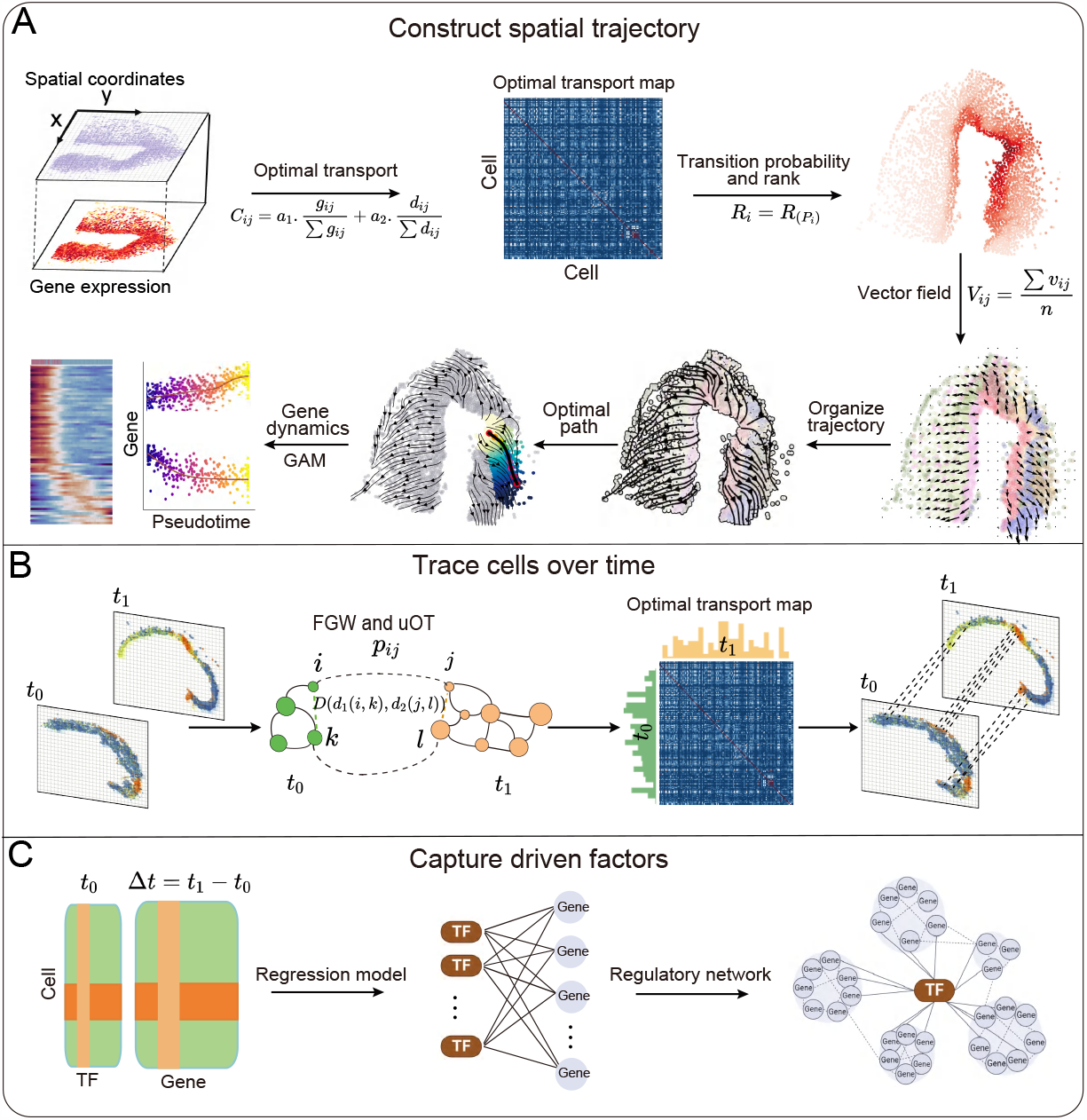
Frameworks of spaTrack. A. Construct cell trajectories from ST data. In brief, we scale the gene expression difference *g*_*ij*_ and spatial distance *d*_*ij*_ to construct the cost matrix of cell transition *C*_*ij*_ . The transition probabilities are estimated by solving the optimal transport problem; Cells are ranked according to their transition probabilities relative to the starting cells; A vector field of cell velocity is built to organize the optimal trajectories; The optimal path between a starting cell and an ending cell is constructed using the least action path method; To identifying pseudotime-dependent genes, we use a generalized additive model (GAM) to fit the dynamics of gene expression along a trajectory. B. Trace cells across multiple samples of a time series. To compute the transport map between cells at time *t*_0_ and *t*_1_, we solve the unbalanced optimal transport (uOT) problem by adapting the Fused Gromov-Wasserstein (FGW) algorithm. *d*_1_(*i, k*), *d*_2_(*j, l*) are the spatial distances between any pair of cells *i, k* at *t*_0_, and their corresponding cells *j, l* at *t*_1_; *D* measures the scaled difference of *d*_1_ and *d*_2_. C. Capture dynamic driven factors. A neural network framework is implanted in the algorithm, with expression profile of genes at *t*_0_ time as input layer, prediction of TF expression of *t*_0_ time as output layer. TF-gene pairs with high weights are screened to build regulatory network.

The cost matrix consists of two components: expression difference and physical distance, both of which employ Euclidean distance as the metric. The algorithm performs dimension reduction from original features and the Euclidean distance in the embedding space is used as the measurement of the difference of expression profile. By appropriately scaling these two distance measurements, we fine-tuned the relative importance of each one by adjusting the weight parameters of *α*_1_ and *α*_2_. The algorithm maximizes the distance of a cell to itself to prevent self-transitions. To solve the OT problem using the cost matrix generated above, an entropic regularization term is introduced. This term helps to produce a smooth probability distribution (Cuturi, 2013). The resulting OT probability matrix displays the likelihood of each cell being transferred to other cells with the minimum cost incorporating both expression profile and spatial coordinates (Figure S1A and S1B). spaTrack makes the assumption that cells with higher transferring probabilities are more closely related during the development. Assigning the starting cells, the algorithm can reorder all other cells according to their transferring probabilities relative to the starting cells (see Methods). Subsequently, spaTrack creates a vector field of the cell velocity averaged from the transferring probabilities and directions of all cells in the neighborhood. Streamlines of the vector field are finally organized and smoothed as the spatial trajectories. Furthermore, we adapted the least action path (LAP) algorithm(Qiu et al., 2022) to construct the optimal path of differentiation between a starting cell and an ending cell. Specifically, the vector field of cell velocity of the transition probability in a spatial neighborhood is computed to replace the RNA-splicing velocity. A set of neighboring cells are mapped to the inferred optimal path to determine cell orders and pseudotimes along the differentiation. The arc length between the mapped anchor point and the starting cell is normalized as the pseudotime of a cell.

Sometimes, ST data from single sample will not capture all cell states of a complete biological process, especially when the process is time coordinated. However, integrating multiple samples will lose the spatial coordinates of each tissue section. spaTrack provides an integrating strategy to accommodate this situation. spaTrack separately computes the transition probability and cell velocity for each ST data; And then integrates the vector fields of all datasets to organize the overall trajectories in an UMAP embedding. This allows for a more accurate and complete representation of the cell trajectories without losing the spatial coordinates of each data.

### Tracing cells across multiple ST data of a time series

Multiple ST data sampled from a time series will provide a spatiotemporal transcriptomic view of biological systems. In order to align cells and generate trajectories across samples of different time intervals, spaTrack adapts an unbalanced and structured OT algorithm considering the uneven expression mass and distributions of samples. ST data of different time intervals are treated as different distributions (Figure 1B). The optimization problem of this OT includes three terms: a measure of the expression profile differences between cells of the two samples; a measure of the spatial distance differences between paired-cells of the two samples; and a measure of the KL divergence (see Methods). spaTrack extends a distance consideration in the optimization problem: If a pair of cells (*i, k*) in time *t*_*i*_ is mapped to a pair of cells (*j, l*) in the next time *t*_*i*+1_ with high probability, the distance between cells *i* and *k* in the time *t*_*i*_ is close to the distance between cells *j* and *l* in the next time *t*_*i+1*_. Therefore, three matrices are included in spaTrack to solve the optimization problem: a gene expression dissimilarity matrix, a spatial distance matrix for cells at time *t*_1_, and a spatial distance matrix for cells at the next time *t*_*i+1*_. The resulting transport plan depicts the transition probabilities of individual cells across ST samples.

After computing transport maps between two adjacent time points, the next step is to extend the transitions to the next time interval. To achieve this, we adopt the Markov assumption that the developmental process follows a memoryless property. Therefore, the long-range transitions could be inferred by composing transport maps using matrix multiplication (see Methods). At each time point, we start from the cells transported from the previous time point and infer their subsequent transitions. This progressive method has the advantage of avoiding direct inference of cell transition over long time intervals, and producing more coherent and credible results. Following this approach, spaTrack is able to establish a long-term and continuous transition map of cell trajectory.

### Modeling dynamic driven factors

spaTrack explores the driven factors regulating cell trajectories, which helps to construct the regulatory network underlying cell differentiation. spaTrack establishes a global regulatory network to interpret the connection between the expression profiles of transcription factors (TFs) at current time and of targeted genes at later time. We propose to set up a regression model to learn the linear continuous function, representing the relationship between TFs and the dynamic change of genes (Figure 1C). A weight matrix is optimized to present the importance of TFs and their genes. This regression model works for both data of discrete time points when the transport map has been constructed, and data of single sample assigned with continuous pseudotime (see Methods). To examine the inferred regulatory network of TFs and targeted genes, we extracted TF-target pairs with top weights and examined their relationship of expression profiles across cells along time. As expected, correlations were observed for those TF-target pairs with top weights (Figure S1C).

### spaTrack constructs reliable spatial trajectories over multiple scenarios

spaTrack provides reliable performance in simulation data where various topologies of differentiation are considered, and quantitative evaluations demonstrate superior performance from spaTrack over existing methods. We designed seven various scenarios in organizing cells temporally and spatially during differentiation (Figure 2A): (1) Continuous differentiation from one core, expanding outwards in an orderly manner; (2) Nonlinear spreading, with faster speed at earlier stage and slowing down later; (3) Fluctuant coordinates, cell expanding to surrounding space with high fluctuation; (4) Discrete differentiation in different niches; (5) Branched lineages in space; (6) Branched lineages without spatial information or of totally infiltrated cells, which is treated as the SC data manner; (7) Multiple samples with time intervals. We applied a lineage-imbedded SC data simulator to generate differentiating cells followed by a spatial assignment according to the seven scenarios (see Methods).

**Figure 2.**
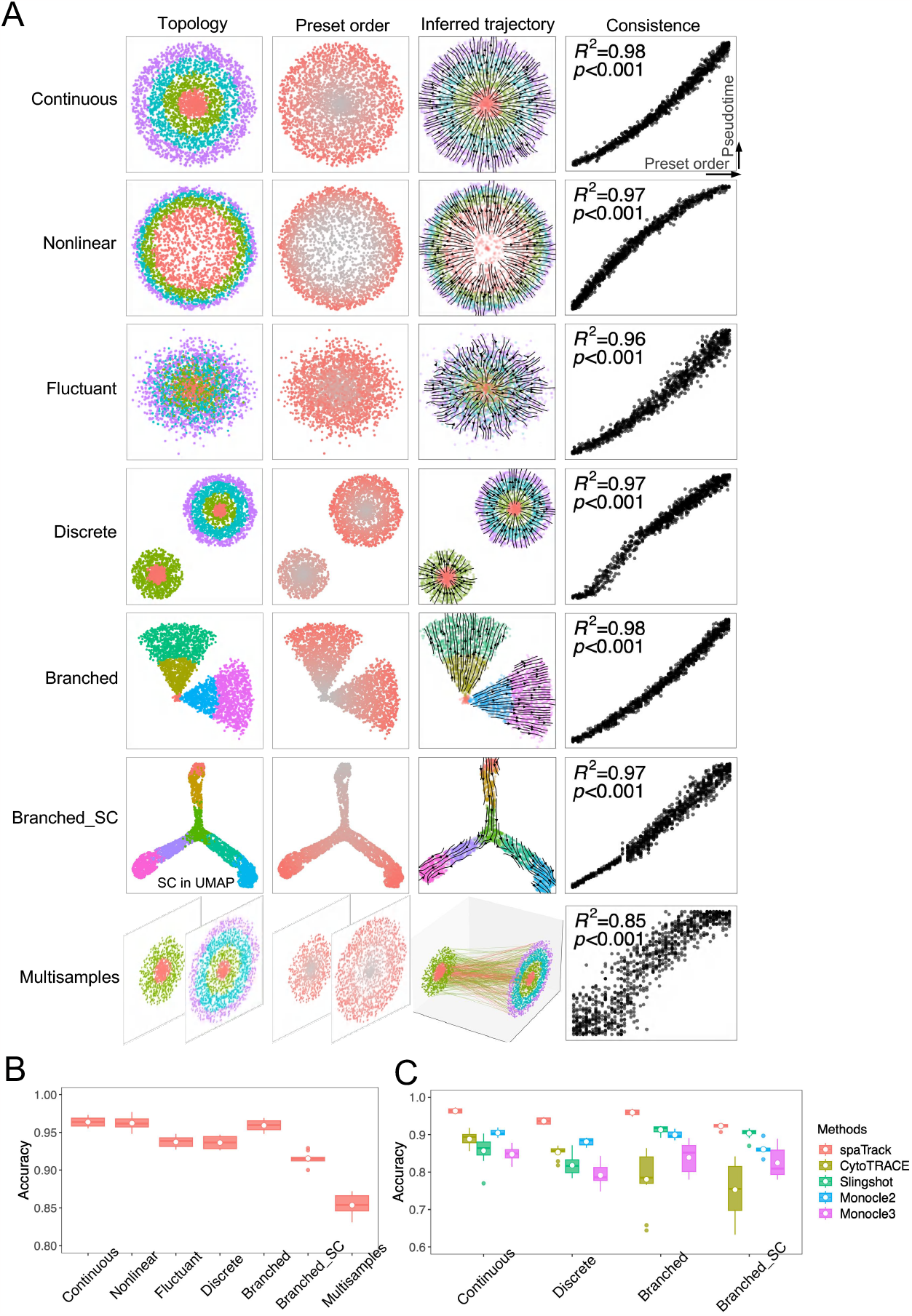
spaTrack constructs reliable spatial trajectories in multiple scenarios. A. spaTrack constructs reliable spatial trajectories in seven scenarios of organizing cells temporally and spatially during differentiation. There are scenarios (rows) of Continuous, Nonlinear, Fluctuant, Discrete, Branched, Branched_SC, and Multisamples. All topologies are spatially organized except for Branched_SC, which is SC data and mapped in UMAP embeddings. The first column is the topology of each scenario in space or UMAP, colors present different cell types; The second column is the heatmap of preset cell orders in simulations; The third column is inferred trajectories from spaTrack; The fourth column is the consistence between inferred pseudotime and preset cell orders, in which cells are randomly sampled from 100 repeats of each scenario. For the spatial scenarios, each ST sample takes a 5000 μm × 5000 μm square in space. B. Accuracy of inferred cell orders in the seven scenarios. The accuracy was calculated from 100 repeats of each scenario. C. Comparison between spaTrack and other commonly used methods applicable for expression matrix.

Based on hundreds of repeats for each scenario, we observed that spaTrack constructs reliable spatial trajectories over multiple scenarios (Figure 2A and 2B). spaTrack presents several advantageous properties in these validations: Firstly, spaTrack captures local details of cell differentiation with spatial trajectories reflecting the true topology. Secondly, spaTrack achieves high consistence with preset cell orders and accuracy even for those scenarios of low space and transcriptome correlation (Nonlinear or Fluctuant scenarios), because both gene expression and spatial coordinates will contribute to the computation of cell transition, making the results robust. Thirdly, for spatially discrete and branched lineages, spaTrack precisely depicts each lineage avoiding interference from each other as the spatial gap depresses their transition probabilities (Discrete or Branched scenarios). Fourthly, spaTrack is totally compatible with SC data when the spatial coordinates are missing or cells are infiltrated without spatial organization (Branched-SC scenario). At last, spaTrack could directly trace cell trajectories of multiple samples of a time series (Multisample scenario), the fine consistence and accuracy making it applicable to trace cells in a wide range of developmental questions such as embryonic development and tumor metastasis, where cells are spatially and temporally organized.

Comparing spaTrack with currently widely-used TI methods for expression data, spaTrack presents superior performance over other methods (Figure 2C). Integrating spatial information adds accuracy of spaTrack comparing with those SC methods without spatial consideration. For the Branched-SC scenario, spaTrack provides closed performance with SC methods since the spatial coordinates are missing. The RNA-velocity based methods are not applicable for the simulated expression matrix. Their performance on ST and SC data will be compared with spaTrack for empirical data in the Discussion section.

According to the highlighted features of the algorithm and the performance evaluated in various scenarios, spaTrack provides several advantageous functions in TI. Firstly, spaTrack could accurately uncover local details of spatial trajectory. Secondly, a single sample may not capture all cell states involved in the complete developmental process. To overcome this limitation, spaTrack could generate and extend the complete trajectories by integrating the transition matrix of multiple samples, without losing their spatial information. Thirdly, direct cell mapping across multiple sections could vividly depict cell trajectories along a time series. Fourthly, spaTrack captures potential driven factors and networks along the time intervals underlying cell differentiation. Furthermore, spaTrack exhibits low resource requirements in terms of both power consumption and computing memory while maintaining a high processing speed (Figure S1D). Specifically, the generation of trajectories using 5k cells with 20,000 features can be accomplished within one minute, while utilizing a modest memory allocation of 6.9 GB.

### spaTrack constructs fine local trajectories of axolotl telencephalon regeneration

We applied spaTrack to reconstruct the spatially detailed trajectories of the regeneration of axolotl telencephalon after injury. Brain regeneration requires the coordination of complex responses in time and region-specific manners. Taking the spatial coordinates of cells into consideration, spaTrack can capture local details of cell trajectory that may be discontinuous in space. Axolotl is a model for studying brain regeneration as its ability to regenerate lost cortical cells after injury. We collected ST data of axolotl samples from a time series after injury(Wei et al., 2022), including samples of 5 days (D5), 10 days (D10), 15 days (D15), and 20 days (D20) (Figure 3A, Figure S2A and S2B), to uncover the regeneration process in space.

**Figure 3.**
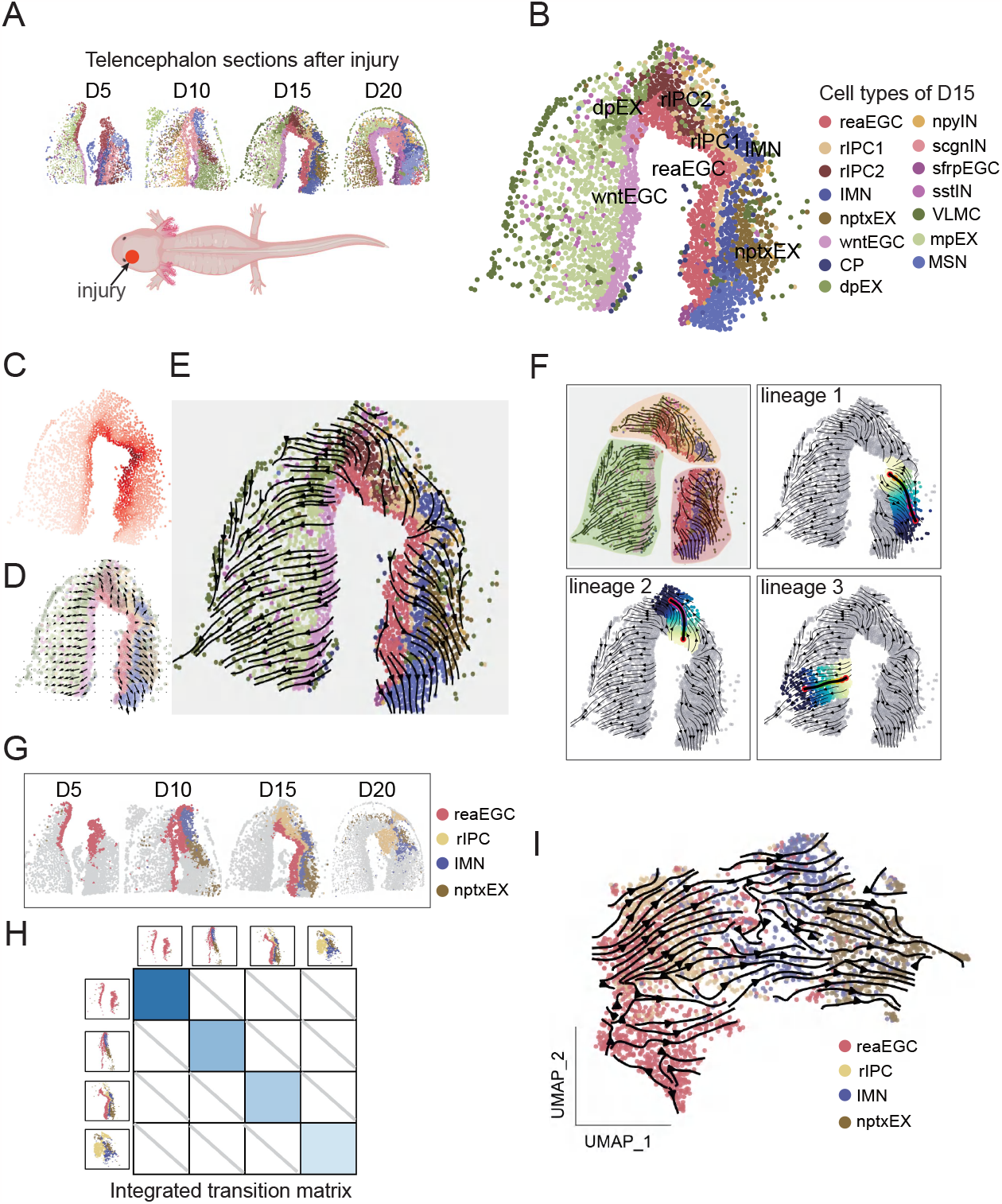
Fine local trajectories of axolotl telencephalon regeneration. A. Collections of the ST data of regenerative stages after injury of axolotl telencephalon at 5 days (D5), 10 days (D10), 15 days (D15), and 20 days (D20). B. Spatial distribution of cell types in regenerative stage of D15. C. Heatmap of transition probability relative to starting cells. D. Vector field of cell velocity, reflecting the direction and potential of transition. E. Regenerative trajectories of D15 inferred by spaTrack. F. Three major lineages of regenerative trajectories, and the optimal path of each lineage. G. Spatial distribution of regeneration-related cell types in the ST data of D5, D10, D15, and D20. H. Integration of transition matrixes from multiple ST samples. I. The complete regenerative trajectories integrating from multiple samples.

D15 shows wound closure with enriched cell types of progenitors and immature neuron cells, including ependymoglial cells (EGC), ractive EGC (reaEGC), immature neuron (IMN), regeneration intermediate progenitor cell (rIPC) and *nptx*^+^ lateral pallium excitatory neuron (nptxEX) (Figure 3B). reaEGC responses to injury and starts the tissue repair suggested by previous studies(Lust et al., 2022; Wei et al., 2022), which presents high proliferative activity (Figure S2C and S2D). Adjacent layers of intermediated cells were observed between reaEGC and nptxEX across the wound area, indicating their transitions during regeneration. spaTrack constructed the local details of regeneration, generating the probability, vector field, and streamlines of cell transition (Figure 3C-3E), uncovering the three spatial lineages of cell differentiation of D15 (Figure 3F). Lineage 1 ranged from wound center to the right-edge of telencephalon, which was reaEGC - rIPC - IMN – nptxEX axis. Lineage 2 was the regeneration of wound area on the dorsal region, which described the complex transitions between reaEGC and IMN, rIPC, and dorsal palliumexcitatory neuron (dpEX). Lineage 3 presented the normal development from *Wnt*^*+*^EGC (wntEGC) to medial pallium excitatory neurons (mpEX). These results depicted the differentiation from reaEGC to intermediate and mature neurons during regeneration after injury, which were consistent with previous reports(Lust et al., 2022; Wei et al., 2022). Importantly, spaTrack explored cell differentiations that were temporally and spatially discontinuous. The trajectory of lineage 2 in the wound area was discrete with the normal trajectory of lineage 3, which were separate processes in development and regeneration.

Single sample presents only a subset of cell types involved in the regeneration process, with sparse cell population and continuity (Figure 3G, Figure S2E). However, integrating all ST samples of D5, D10, D15 and D20 will lose the spatial coordinates of each axolotl tissue section. To address this issue, spaTrack implements an integrating framework to separately calculate cell-transition probability of each sample, and next integrate all transition matrix for the inference of complete trajectory. From an integrated probability matrix (Figure 3H), spaTrack generated the complete trajectories of regeneration and visualized on their UMAP embeddings (Figure 3I). Abundant intermediated cells rIPC and IMN were fully captured showing better continuity than only one sample (Figure S2E and S2F).

### Tracing neuron cells across mouse embryos of a time series

spaTrack provides a novel strategy to trace cells across multiple ST samples by direct mapping cells via an unbalanced OT strategy. Development of mouse embryos requires strict spatial-temporal organization. To probe the dynamics of early neurogenesis, we applied spaTrack on the ST data of developing dorsal midbrain of mouse embryos at day 12.5 (E12.5), 14.5 (E14.5), and 16.5 (E16.5) (Figure 4A, Figure S3A and S3B). Radial glia cells (RGC) are reported as the progenitors of both neuroblasts (NeuB) and glioblasts (GlioB)(Chen et al., 2022a), but their spatial-temporal transitions are not well characterized. RGC decreases from E12.5 to E14.5 and E16.5, while NeuB and GlioB expand in E14.5 and E16.5, and are not evenly distributed along the spatial axis (Figure 4A). spaTrack optimally transported cells from E12.5 to E14.5, and subsequently to E16.5 (Figure 4B), tracing their dynamic differentiation across time. At E14.5, 81% of the successfully transported NeuB cells were found originating from RGC of E12.5, while the corresponding number for GlioB was 74%. Subsequently, at E16.5, 51% of the successfully transported NeuB and 42% of GlioB were transported from RGC of stage E14.5, indicating the RGC is the main source of NeuB and GlioB (Figure 4B). Visualizing these mappings in space, we directly observed the coordinately organization of differentiation in each time point (Figure 4C and 4D, Figure S3E). Differentiations of RGC-NeuB and RGC-GlioB were restricted to different regions and embryonic stages. RGC in the rostral axis mainly differentiated into NeuB, and RGC in the dorsal and caudal regions differentiated into GlioB. RGC-NeuB differentiation mainly occurred from E12.5-E14.5, while RGC-GlioB arose between E14.5-E16.5. All of these results of spaTrack suggested neurogenesis and gliogenesis were asynchronous and spatially heterogeneous, consistent with previous findings (Chen et al., 2022a).

**Figure 4.**
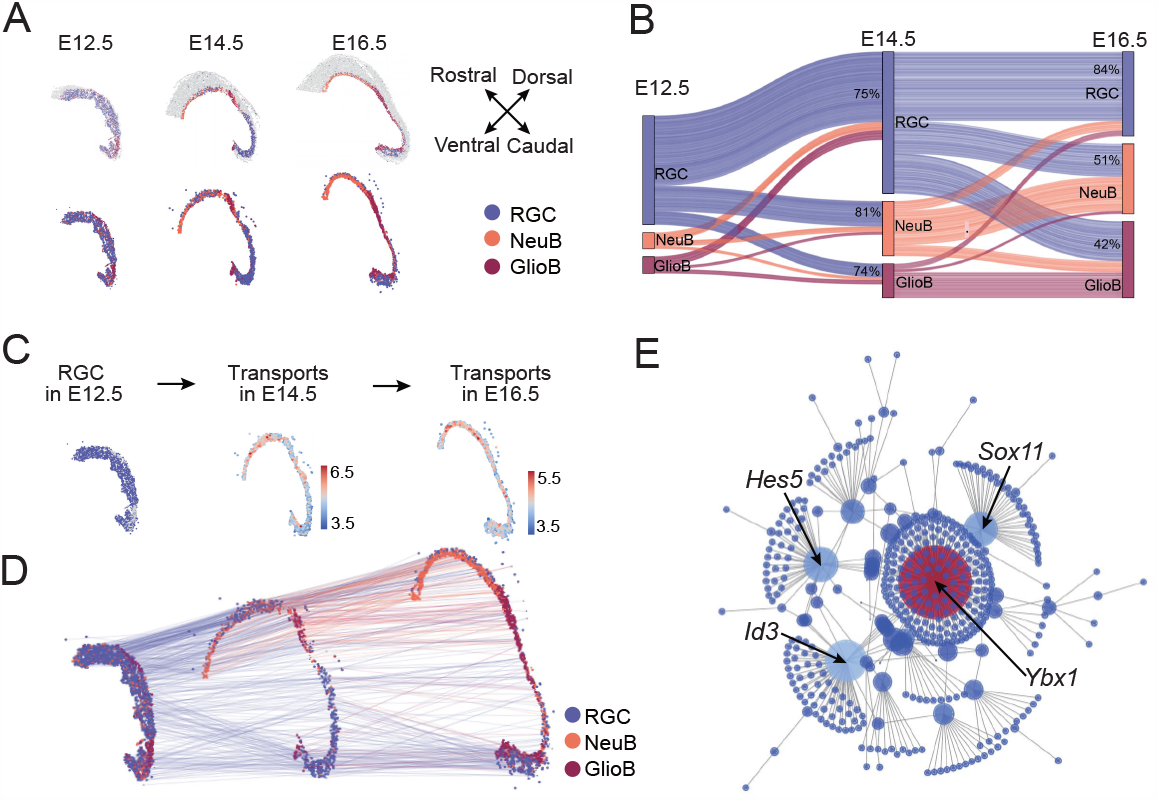
Tracing neuron cells across mouse embryos of a time series. A. ST data of dorsal midbrain regions of mouse embryo at day 12.5 (E12.5), 14.5 (E14.5), and 16.5 (E16.5). Spatial distribution of radial glia cells (RGC), neuroblasts (NeuB), and glioblasts (GlioB) are plotted. B. Sankey diagram of cell tracing across temporal sections. Blue segment represents RGC, brown for NeuB, red for GlioB. Percentage of RGC-derived cells in all successfully transported cells of each type is labeled. C. Tracing the transition of RGC in E12.5 (left) to E14.5 (middle) and E16.5 (right). Probabilities of successfully transported cells are plotted. D. Visualization of the transition trajectories of RGC across samples of different time. E. Regulatory network underlying the RGC differentiation over time.

Furthermore, driven factors of the neuron differentiation were investigated by a regression model in spaTrack. A regulatory network was built to present the connection between TFs and targets along the temporal intervals (Figure 4E). Several TFs were highlighted by our method. *Ybx1* reported as a crucial factor for forebrain specification and restricting mid-hindbrain growth in mouse embryo, fine-tunes the spatiotemporal expression of neurodevelopmental genes (Evans et al., 2020). Another TF *Sox11* is required in embryonic neurogenesis and *Sox11*-depleted embryos develop small and disorganized brains, accompanied by transient deficits in neural progenitor cells (Wang et al., 2013). Therefore, spaTrack could provide valuable reference and methodological support for the advancement of the neuroscience field.

### Recovering the diverse trajectories of tumor expansion

Intratumoral heterogeneity manifests as spatial heterogeneity, which describes the uneven distribution of diverse malignant subclones within tumor, and as temporal heterogeneity, referring dynamic variations in tumor populations and molecules over time (Dagogo-Jack and Shaw, 2018; Hausser and Alon, 2020). Tumor heterogeneity drives tumor progress and drug resistance, creating the need to quantitatively investigate tumor subclones and dynamics over space and time.

We collected ST data from a primary tumor section of intrahepatic cholangiocarcinoma (ICC) (Wu et al., 2023), covering the regions of both intratumor and boundary (Figure 5A, Figure S4A-S4C). Previous study(Wu et al., 2023) detected strong immunosuppression and metabolic reprogramming in the invasive zone of tumor boundary, suggesting the spatially diversity of tumor progress. Eight subclones were identified in the primary ICC tumor (P0-P7) (Figure 5B), with P0 showed pronounced expression of stemness markers and cell cycling genes (Figure 5C and 5D, Figure S4D). Applying spaTrack to reconstruct cell trajectory of malignant cells assigning P0 as starting cells (Figure 5E), we identified three diverse lineages starting from P0 and spanning the tumor space in three directions (Figure 5E and 5F). Lineage 1 (P0-P1-P2-P7) expanded to the border region between tumor and hepatic cells. Lineage 2 (P0-P3-P4) extended to the tumor bottom and lineage 3 (P0-P5-P6) elongated along the top area. To determine biological difference among the three lineages, we identified trajectory-depended genes by fitting a generalized additive model between the pseudotime and gene expression along the optimal path, inferred using the LAP method (Figure 5G and 5H, Figure S4E and S4F). In the associated genes of lineage 1, *COL1A1* is a major component of the tumor extracellular matrix related with tumor development and immune profile(Chen et al., 2022b). *SAA1* and *SAA2* lead to recruitment and polarization of macrophages, promoting local immunosuppression (Wu et al., 2023). Annotating associated genes of lineage 1, we observed significant enrichment of ECM organization and regulation of platelet and neutrophils (Figure 5I), which involve in tumor migration, metastasis, and immunosuppression(Winkler et al., 2020; Wu et al., 2023). The GSEA scores of ECM and EMT pathways further indicated the metastatic potential of lineage 1 (Figure 5J and 5K). Additionally, spaTrack constructed the regulatory network underlying lineage 1 (Figure S4G), capturing the TFs and targets of tumor growth and metastasis, e.g. *KLF7* (Gupta et al., 2020) and *ETS2* (Zhang et al., 2021). All these characters were not observed in the other two lineages, indicating the spatial heterogeneity of tumor progress.

**Figure 5.**
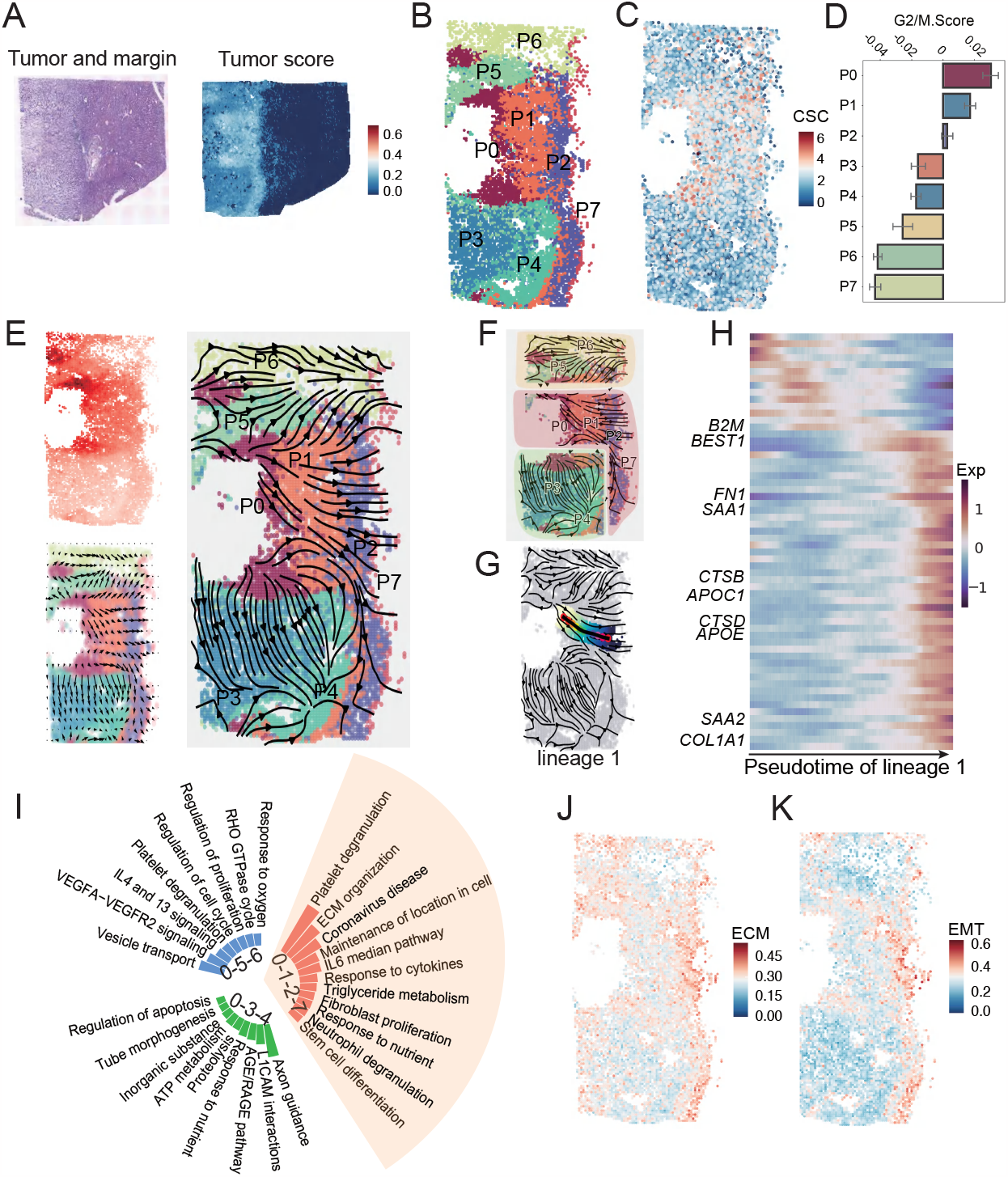
Recovering the diverse trajectories of tumor expansion. A. A ST sample of primary tumor of intrahepatic cholangiocarcinoma (ICC). H&E staining (left), and distribution of malignant cells (right) are plotted. B. Spatial distribution of tumor subclones. C. Spatial expression of cancer stem cell markers. Expression of gene CD44, ID1, CDH1, and FOSL1 are summed up. D. The G2M score of tumor subclones. E. Trajectories of tumor expansion. The transition probability relative to starting cells (left top) and vector field of transition velocity (left bottom) are visualized. F. Three lineages of tumor trajectories in space. G. The optimal path of the lineage 1 (P0-P1-P2-P7). H. Pseudotime-dependent genes of lineage 1, screened by fitting a generalized additive model. Only top 10 significant genes are labeled. I. Functional annotation of pseudotime-dependent genes of the three lineages. J. Gene score of ECM pathway. Gene score is calculated as the averaged expression of the genes in each pathway. K. Gene score of EMT pathway.

### Tracing tumor metastasis

spaTrack provides the ability to trace cells across tissues of different time/conditions and therefore could reconstruct the trajectory of tumor metastasis. Tumor metastasis refers to the process by which cancer cells detach from the primary tumor and spread through the bloodstream or lymphatic system to colonize distant organs (de Visser and Joyce, 2023). Understanding the origins and colonizing process of tumor metastasis, provides important insights in developing effective strategies to target metastatic relapse and improve patient outcomes (Ganesh and Massague, 2021).

We collected a metastatic tumor (Figure 6A, Figure S5A and S5B) from the lymph node corresponding with the primary ICC tumor (Figure 5A) in the same patient (Wu et al., 2023). Malignant cells at the metastatic site were categorized into four major clusters (M0-M3) (Figure 6B), forming a layered structure. M0 located at the core site of the tumor; M1 lied in the middle layer; M2 and M3 covered the outer layer. We applied spaTrack to optimally transport the malignant cells from the primary tumor to the metastatic tumor. M3 showed numerous successful transports from P0/P1, which was significantly higher than any other pairs of clusters (Figure 6C). By plotting the successful transports between the primary and metastatic tumors (Figure 6D), we observed malignant cells of the primary tumor (mainly from P0-P1) implanted to the bottom axis of the metastatic tumor, belonging to subclone M3, from where the metastatic cells putatively expanded to a new tumor. To further investigate the origins and colonization of metastatic cells, we inferred and compared the genetic variants (SNP) of the malignant cells from both tumors. P0-P1 shares more variants with M3, than any other pairs of subclones after adjusting the population size (Figure 6E), confirming the metastatic connection inferred by spaTrack. Furthermore, integrating the ST data of the primary and metastatic tumors in the SC manner, M3 approximated with P0 and P1 in the UMAP embedding space (Figure S5C), which was consistent with the results of spaTrack. Constructing the regulatory network between the primary tumor and the metastatic tumor, we observed HMGA1, ID2, and CEBPG as the key factors driving the metastatic dynamics, all of which play important roles in tumor progression and metastasis (Huang et al., 2020; Sgubin et al., 2022; Sikder et al., 2003) (Figure 6F).

**Figure 6.**
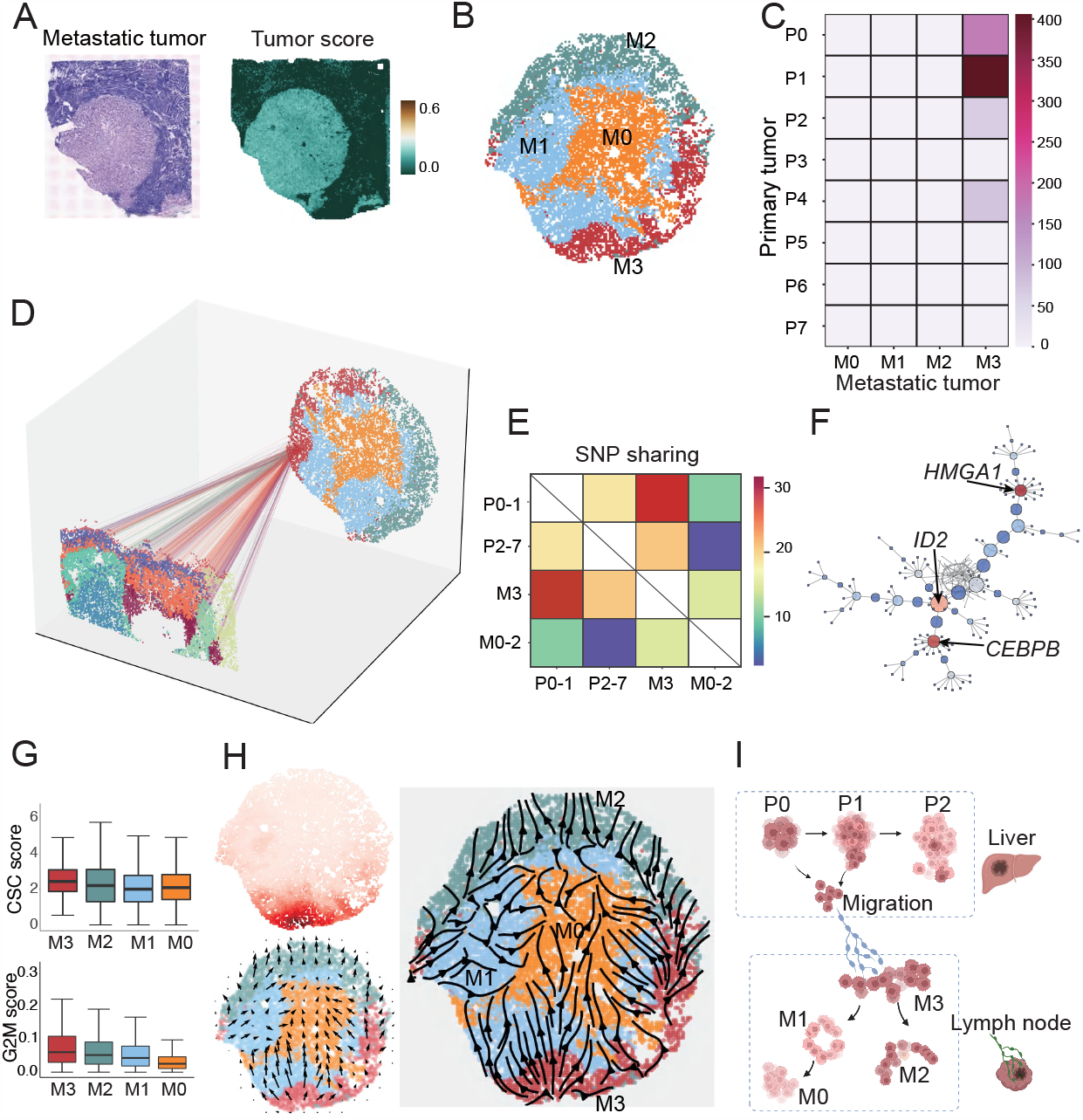
Tracing tumor metastasis. A. A ST sample of metastatic tumor in lymph node, corresponding with the primary tumor of ICC in Figure 5A. H&E staining (left) of tissue, and distribution of malignant cells (right) are plotted. B. Spatial distribution of subclones in the metastatic tumor. C. Counts of optimally transported cells from primary tumor to metastatic tumor. The successful transports were determined by their transition probability. D. Tracing the successful transports between primary tumor and metastatic tumor. E. Expression variants (SNP) shared between primary tumor and metastatic tumor. The number of shared variants is adjusted by population size of clusters. The sharing numbers are significantly higher between P0-P1 and M3 than any other pair of clusters (p<0.05 in one sample t-test). F. Regulatory network underlying the metastasis between the two tumors. G. The expression of stemness markers and G2M scores in subclones of metastatic tumor. H. Trajectories of tumor colonization in the metastatic site. The transition probability relative to start cells (left top) and vector field of transition velocity (left bottom) are visualized. I. A model of the tumor metastasis uncovered by spaTrack.

We subsequently examined the stemness and cell cycling of metastatic cells, both of which indicated M3 presenting the activation of proliferation and expansion in the metastatic site (Figure 6G, Figure S5D and S5E). We applied spaTrack to generate trajectories of the metastatic cells, assigning the successfully transported cells of M3 as starting cells (Figure 6H). It appeared that M3 initiated the colonization along the bottom axis and subsequently progressed to M2 where they formed the outer layer. Both M3 and M2 expanded towards the middle layer (M1) and formed the core site. Overall, spaTrack vividly described the dynamic process of tumor metastasis including origination, colonization, and expansion (Figure 6I). This comprehensive analysis would certainly provide us valuable insights of tumor metastasis.

## Discussion

Development of biological systems commonly requires strict spatial-temporal organization. Spatial coordinates and experimental time might be leveraged as important constraints supplementing to transcriptomic profiles in the TI work. spaTrack, presented as an innovative algorithm that uses the mathematical concept of OT, generates accurate and informative spatial trajectories by incorporating both gene expression profiles and spatial-temporal information from ST data. According to the highlighted features of the algorithm (Figure 1A-1C), spaTrack could (1) reconstruct fine local trajectory from ST data; (2) integrate spatial transition matrix of multiple samples to generate complete trajectories; (3) trace cell trajectory across temporal samples via direct OT mappings; (4) capture developmental driven factors by modelling a function of predicting gene profile at later time by TF expression at current time.

spaTrack has been undergone extensive testing on both ST data and SC data of simulated scenarios, comparing with currently widely used TI methods for expression data (Fig 2A-2C). OT framework has the natural advantage of incorporating spatial distance into the cost measure of cell transition and therefore captures local details and generate spatial disconnected trajectories. Moreover, to compare spaTrack with RNA-velocity based methods, we applied spaTrack, scVelo, and Dynamo, which could directly generate trajectories in spatial coordinates, on the ST data of axolotl telencephalon regeneration (Figure S6A and S6B), which is a comprehensively studied model. As we described before, spaTrack uncovered three spatial trajectories reflecting the true regenerative process. Regeneration trajectories in the wound area are disconnected with those in normal tissue. scVelo presented confusing trajectories with multiple starting spots, which could not be adjusted by simply reversing the velocity direction. Dynamo performed well in most regions, but showed continuity between lineages of temporally disconnected. Comparatively, Monocle3, showed a skeleton along the data shape, without single cell trajectories (Figure S6B), which is a typical result of SC methods using expression data.

Furthermore, we tested and compared the performance of spaTrack and other methods on a complex topology of SC data of primary human hematopoietic stem and progenitor cells (HSPCs) (Qin et al., 2021) (Figure S6C). Human hematopoiesis is a continuously hierarchical process and is comprehensively investigated by previous studies (Buenrostro et al., 2018; Ranzoni et al., 2021). The development of HSPCs follows a branched structure with HSC as the root. spaTrack successfully reconstructed the trajectories that closely recapitulate the established knowledge of hematopoiesis (Figure S6D). In comparison, scVelo generated several nonsensical reversing trajectories, starting from intermediated cell types of active cell-cycling states (e.g., erythrocyte progenitor (pro-ery), B cell progenitors (proB), and granulocyte and monocyte progenitor (GMP)). Without metabolic labels, Dynamo also showed reverse streamlines in erythrocyte and myeloid trajectories. Monocle3 generated a proper skeleton of hematopoiesis but missing the single cell details.

At last, spaTrack requires feasible computing power and memory (Figure S1D), making it a fast and effective option for TI study of ST data. Under a standard CPU thread (Intel(R) Xeon(R) CPU E5-2650 v4 @ 2.20GHz), spaTrack requires only minutes to finish the computation of 5k – 400k cells (with 20,000 features). The memory load depends seriously on the population size, which follows an exponential growth with 6.9 GB for 5k cells.

## ACKNOWLEDGMENTS

This study was supported by funding from the National Key Research and Development Program of China (No.2022YFA1103400 and 2021YFC2501900), and the Guangdong Basic and Applied Basic Research Foundation (No. 2021A1515110832). This work was also supported by the China National GeneBank (CNGB).

## AUTHOR CONTRIBUTIONS

P.F.Q., H.M.Z., L.W., and X.X. designed the study; X.N.S., L.L.Z., Z.F.Y., H.M.Z., and P.F.Q. developed the methodology; Z.F.Y., L.L.Z., X.N.S., and K.H. developed the software and online tutorials; K.H., P.F.Q., L.L.Z., and Z.F.Y. performed the data analysis; X.X.Z., Q.C.Y., Z.Y.L., L.W., and X.Y.W. assisted with data collection and analysis; P.F.Q., X.N.S., K.H., L.L.Z., and Z.F.Y. wrote the manuscript; P.F.Q., H.M.Z., L.W., X.X., and X.J. supervised the study.

## Data resources

The public ST data and SC data used in this study were collected as follows: ST data of Stereo-seq of axolotl telencephalon after injury was obtained from China National GeneBank DataBase (CNGBdb) with accession number CNP0002068. We selected four samples of 5, 10, 15, and 20 days after injury. ST data (Stereo-seq) of mouse midbrain development were collected from CNGBdb with accession number CNP0001543. Three ST samples (Stereo-seq) of mouse embryo sections at 12.5 day, 14.5 day, and 16.5 day were downloaded from CNGB with accession number CNP0002199, including one primary tumor of intrahepatic cholangiocarcinoma (ICC), and one corresponding metastatic tumor. One SC sample (10x genomics) of human hematopoietic stem and progenitor cells (HSPCs) were downloaded from the Genome Sequence Archive of CNCB-NGDC (National Genomics Data Center of China National Center for Bioinformation), with accession number HRA000084.

## Code availability

The open-source software spaTrack is available at https://github.com/yzf072/spaTrack. The tutorial of spaTrack is deposited at https://spatrack.readthedocs.io/en/latest/index.html.

## Methods

### Inferring cell trajectories from single ST data

#### Construction of cost matrix

To reduce computational burden, we perform Principal Component Analysis (PCA) to reduce the dimensionality of the data. Subsequently, we select the top 10 PCA components (defaulting to 30) for downstream analysis. To construct the cost matrix, we incorporate both gene expression profiles and physical distances. Differences in gene expression profile and cell coordinates are quantified using Euclidean distance. For cell i and cell j, the Euclidean distances of gene expression (*g*_*ij*_) and physical distance (*d*_*ij*_) are calculated as follows:

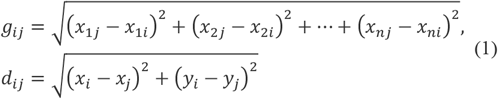

Where n represents the number of principal components selected in the previous step of PCA.

To balance the contributions of the two distance measurements, we first normalize the distances. We then integrate the normalized gene expression distance *g*_*ij*_ and normalize spatial distance *d*_*ij*_ by scaling factors *α*_1_ and *α*_2_ to compute the cost matrix *C*_*ij*_ of cell transition. These factors control the relative importance of each distance measurement, with suggesting values for *α*_1_ and *α*_2_ are between 0 and 1. To prevent self-transitions, the cost matrix is re-defined. When *i* = *j*, we set the cost to the maximum of *C*_*ij*_ times 10^7^, so that the cost of a self-transition is maximized:

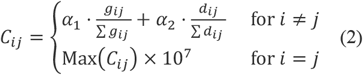

#### Transition probability between cells

Adapting the concept of optimal transport (OT), we calculate the transition matrix by solving the following optimization problem:

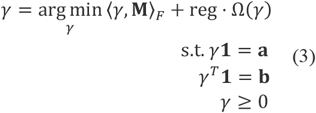

Where **M** is the cost matrix calculated above, Ω is the entropic regularization term Ω(γ) = ∑_*i,j*_ γ_*i,j*_ log.(γ_*i,j*_), **a** and **b** are source and target weights (both sum to 1).

#### Cell order assignment

To utilize spaTrack for cell trajectory analysis, we specify the starting cells as ancestral cells. This can be achieved through various means including importing cell coordinates, cell type, or by manual selection using the interactive user interface to create lasso spots. Once the starting cells are defined, spaTrack will assign cell orders or directions relative to the starting cells for other cells in the dataset. This can be achieved by calculating the transferring probability of each cell to the starting cells. The probability can be determined by summing the transition probabilities of the cell to each of the starting cells.

Let cell_1_, cell_2_, … cell_s_ as starting spots, for cell_i_, the sum of transition probabilities is calculated as:

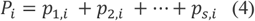

Where the *P*_*i*_ is the probability of starting spots transferred to cell i. We assign a cell order to each cell by assuming that cells with a higher probability of transferring to their ancestors are closer to the ancestors in the trajectory. The probabilities of starting cells transferring to each cell were ranked ascending as:

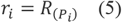

Assuming the same interval, cell orders are normalized using the following formula:

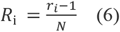

Where N is the total cell number.

#### Cell velocity and organizing trajectory

Cell velocity is defined as the overall transition probability and direction from a cell to its neighbors. Before calculating cell velocity, the neighboring cells are determined using their spatial coordinates and their cell PCA embedding matrix. Users are allowed to choose the number of neighboring cells to consider. The spatial neighbors are identified using the K nearest neighbors (KNN) algorithm, while the PCA matrix neighbors are determined using Euclidean distance between cells. The final set of neighboring cells is obtained by taking the intersection of the two sets of neighbors.

For each cell i, the n neighbors are selected. The velocity between cell i and cell j (j ≤ n) is defined as following:

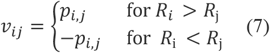

Then the final velocity of cell i is calculated by averaging the velocities of cell i in its neighborhood:

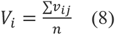

The trajectory is organized from the vector field of cell velocities, which is adapted from sctour ^19^. Briefly, the optimal transition probability matrix is used as weights to calculate the unitary displacement vector for each cell. Only n KNNs of each cell are considered (n = [total spot number/50]):

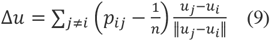

Where *u*_*i*_ and *u*_*j*_ were the coordinates of cell i and j.

#### Optimal path between two cells and pseudotime calculation

To study the differentiation trajectory between two cells over space, we adapt the least action path (LAP) algorithm (Qiu et al., 2022) to construct the optimal path between a starting spot and ending spot. Firstly, we construct a vector field of cell velocity from transition probability instead of RNA velocity, as described in *Formula 7* and *8*, which enables the estimation of cell velocity at any coordinate point. Secondly, given a starting cell and an ending cell, the initial path will be a line connecting the two points. The path is adjusted according to the cell velocity following the LAP algorithm. We will get an optimal path that best fits the transition between the two cells. Afterwards, we need to map all the cells around the optimal path to assign cell orders and pseudotimes along the differentiation. We use the k-Nearest Neighbor (KNN) method to search cells spatially around the path. The neighboring cells are vertically mapped to the optimal path, and the order of the cells is determined according to the mapped anchor point relative to the starting cell. Pseudotimes are defined as arc length between the mapped anchor point and the starting cell. Pseudotimes are normalized to a 0-1 range by dividing the total length of the path.

### Tracing cells across multiple ST data with time-intervals

#### Unbalanced transport across multiple ST data

To compute the transport map between cells at time *t*_*i*_ and *t*_*i*<1_, assuming that there are *m* cells at time *t*_*i*_ and *n* cells at time *t*_*i*<1_, we solve the following optimization problem:

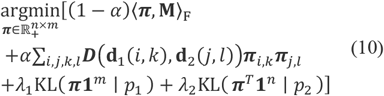

Where **M** ∈ ℝ^n×m^ measures the gene expression dissimilarity between cells of two samples, and ⟨*A, B*⟩_F_ denotes Frobenius inner product of matrices *A* and *B*, and d_1_(*i, k*), d_2_(*j, l*) are the spatial distances between cells *i, k* and their corresponding cells *j, l* at different times respectively, and ***D*** measures the difference between scaled distances (Euclidean norm ∥ ⋅∥^2^). In addition, λ_1_ and λ_2_ are regularization parameters and *p*_1_ and *p*_2_ are weight vectors of each cell. By default, 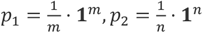,where**1**^*m*^, **1**^*n*^denotes a column vector of length *m, n* containing all ones. The transport problem is solved with following considerations:

1. If cell *i* in time *t*_*i*_ is mapped to cell *j* in the next time *t*_*i*<1_ with a high weight **π**_*ij*_, then the expression profile *x*_*i*_ of cell *i* is similar to the expression profile *x*_*j*_ of cell *j*.
2. If a pair of cells (*i, k*) in time *t*_*i*_ is mapped to a pair of cells (*j, l*) in the next time *t*_*i*<1_ with high weights **π**_*ij*_ and **π**_*KI*_, then the distance ***d***_**1**_(*i, k*) between cells *i* and *k* in the first time *t*_*i*_ is close to the distance ***d***_***2***_(*j, l*) between cells *j* and *l* in the next time *t*_*i*<1_.
3. Unbalanced optimal transport (Chizat et al., 2018), is with a more realistic approach to solving practical problems, for instance, it is suitable for scenarios where batch effects are present at different time points or when investigating the impact of the varying numbers of cells with value-added differentiation.

The sum of the first two terms in *Formula 10* represents a classic Fused Gromov-Wasserstein algorithm(Titouan et al., 2019). By introducing the last term, we extend the structured transport to handle unbalanced transport problems, where the equality constraints are relaxed to impose bounds on the marginals of the transport plan using of KL-divergence measure.

#### Computing trajectories of interest cells

At a given time point, a collection of starting cells can represent a specific cell type or any region of interest in space. Then the distribution of descendant cells at the next time point *t*_*i*<1_ can be calculated based on the transition matrix,

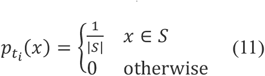

In which S is the set of starting cells. The descendant distribution can be calculated as following

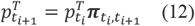

where 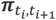 is the optimal transport map between *t*_*i*_ and *t*_*i*<1_ calculating from (10).

### Learning gene regulatory models

OT has the capability to capture potential driven dynamics. We interpret the vector field as a model of gene regulation, which establishes functional relationships between the expression of transcription factors (TFs) at current time point and the expression of genes in a period of time. We propose to set up a regression model to learn the positive/negative regulation of genes by TFs. For ST data of two time points, we sample pairs of cells with expression (*X*_*i*_, *X*_*i*<1_) from the transport map and calculate gene changes Δ_*i*_:

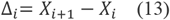

Then, we extract TF expression from time *i* + 1 and construct the following regression:

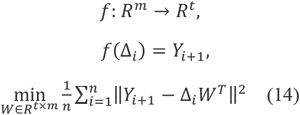

where *Y*_*i*+1_ = *X*_*i*+1_*T, X*_*i*_ ∈ *R*^1×*m*^, *X*_*i*+1_ ∈ *R*^1×*m*^ denotes the gene expression of pairwise mapped cells at two time points with m genes, *f* is the learned linear continuous function, representing the relationship between genes and TFs, *W* ∈ *R*^*t*×*m*^ denotes weight matrix with TFs and genes, *T* ∈ *m* × *t* stands for one-hot encoding matrix of TFs and genes. Here, we perform min-max normalization for gene changes and TF expression respectively and put data into the regression model to get weights.

For ST data of a single tissue section, we format the data to adapt the regression model. Cells are sorted according to their inferred pseudotime and are averagely grouped according to the setting bins. Then, cells of each pair of adjacent bins could be inputted to the model, which are processed in the same way.

To solve the problems of sparse data and reduce slow convergence, we used a meta-analysis method. We repeatedly select random cells and calculate the mean expression as new data, to improve data quality and increase sample size. To avoid the instability caused by random initialization of the model, we take the average of ten random training as the final result. Finally, top positive/negative correlation weight pairs are sorted from high to low and stored in data frame format. The regulatory network of TFs and genes can be displayed visually.

### Simulations of ST data

We applied a lineage-imbedded SC data simulator, Splatter (Zappia et al., 2017), to generate differentiating cells followed by a spatial assignment according to various scenarios. Basic parameters to restrain the expression of single cells in the simulator were assigned as nGenes=3000, batchCells=3000, mean.shape=0.6, mean.rate=0.3, bcv.common=0.2, dropout.mid=0, dropout.shape=-1, out.prob=0.05, de.prob=1; Lineage parameters were assigned according to the topologies: method = “paths”, path.length= from 60 to 100, path.skew= from 0.2 to 0.5, path.nonlinearProb=0.1. After the simulations of expression matrix of SC, we assigned the 3000 cells of each simulation to a 5000 μm × 5000 μm square, assuming each cell taking up a 50 μm × 50 μm spot. For a spatial assignment, cells were organized according to their preset steps expanding from the center, with a fluctuation of a normal distribution *µ* = 0, *σ* = 3 *to* 6 . For each scenario, simulations were repeated 100 times. Consistence and accuracy were evaluated from all these batches. Accuracy was estimated as the fraction of consistent cell orders of any random cell pairs compared to the preset orders.

### Processing of raw data of ICC and its metastatic tumor

We downloaded the Stereo-seq GEM file of the primary tumor of ICC and its metastatic tumor from a previous study^22^. The GEM file includes the DNB coordinates and gene UMI counts in each DNB (220 nm). It was difficult to segment cells of tumor tissue and assemble the reads of single cells. We therefore merged 100 × 100 DNBs into a single informative ‘bin’ as a pseudo-cell (50 μm × 50 μm in square). To remove low-quality data, cells with expressed genes number < 500, expressed genes UMI <500 and a proportion of mitochondrial UMI > 20% were removed from downstream analysis. Finally, we obtained a total of 19908 and 28609 cells for the primary tumor and metastatic tumor respectively. The quality details of the data showed by violin and heatmap plots were presented in Figure S4A and S4B, Figure S5A and S5B.

### Cell type deconvolution of ST data and identification of malignant cells

ST data were deconvoluted using the seeded NMF method implemented in SPOTlight v0.17 ^27^. SC data were used as references to infer the composition of each ST bin ^22^. A default threshold of 0.08 was applied to filter the composition of cell type. The distribution of malignant cells was further examined by marker genes. We removed bins with high expression of marker genes relating with T cell, B cell, macrophage, fibroblast and endothelia cells. Finally, we obtained a total of 6,470 and 7,927 malignant cells of the primary tumor and metastatic tumor respectively. BayesSpace^28^ was performed to cluster cells with spatial coordinate.

### Identifying the genetic variants in ST data

In order to reliably detect single-cell expressed variants, we pooled all reads of tumor cells together to call variants (SNP). Tumor cells were determined by annotation of SPOTlight. Both Samtools (Li et al., 2009)(v1.16) and Strelka (Saunders et al., 2012) are applied to call the variants. Successful callings from both methods were used for downstream analysis. Due to the sparsity of the ST data, we used following criteria of filtration to reduce the artifacts and false positives: variants covered by at least 70 reads; reads with alternative variant take up >5% of all reads; variants observed in at least 3 tumor cells.

Shared SNPs between clusters of the primary tumor and the metastatic tumor were calculated. To yield more dependable comparisons, we performed 30 repetitions of counting of shared SNPs by random cell sampling from clusters. For each counting and comparison, cells of clusters were sampled to equal population size.

### Identifying pseudotime-dependent genes

spaTrack applies generalized additive model to fit the dynamics of gene expression along a trajectory. For each gene, spaTrack fits the expression changes and the corresponding pseudotime value of cells using the generalized additive model in pyGAM package. The formula of the model is as:

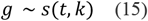

Where *g* represents the gene expression in cells; *t* denotes pseudotime value of all cells along a trajectory; The function k is a spline function used as a piecewise polynomial to fit smooth curves. P-values are adjusted for multiple testing using the BH method.

To determine whether the dynamics of gene expression across trajectory is decreasing or increasing, spaTrack calculates JS score between actual expression and standard downward/upward trends using following formula:

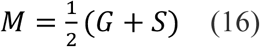

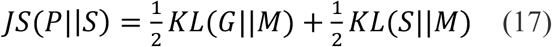

Where, G represents predicted gene expression from the model, S represents a set of standard downward-or upward-trend values. KL is calculated by python SciPy packages.

## Reference

Bergen, V., Lange, M., Peidli, S., Wolf, F.A., and Theis, F.J.J.N.b. (2020). Generalizing RNA velocity to transient cell states through dynamical modeling. 38, 1408–1414.

Buenrostro, J.D., Corces, M.R., Lareau, C.A., Wu, B., Schep, A.N., Aryee, M.J., Majeti, R., Chang, H.Y., and Greenleaf, W.J.J.C. (2018). Integrated single-cell analysis maps the continuous regulatory landscape of human hematopoietic differentiation. 173, 1535–1548. e1516.

Cang, Z., and Nie, Q.J.N.c. (2020). Inferring spatial and signaling relationships between cells from single cell transcriptomic data. 11, 2084.

Cang, Z., Zhao, Y., Almet, A.A., Stabell, A., Ramos, R., Plikus, M.V., Atwood, S.X., and Nie, Q.J.N.M. (2023). Screening cell–cell communication in spatial transcriptomics via collective optimal transport. 20, 218–228.

Cao, J., Spielmann, M., Qiu, X., Huang, X., Ibrahim, D.M., Hill, A.J., Zhang, F., Mundlos, S., Christiansen, L., and Steemers, F.J.J.N. (2019). The single-cell transcriptional landscape of mammalian organogenesis. 566, 496–502.

Chen, A., Liao, S., Cheng, M., Ma, K., Wu, L., Lai, Y., Qiu, X., Yang, J., Xu, J., and Hao, S.J.C. (2022a). Spatiotemporal transcriptomic atlas of mouse organogenesis using DNA nanoball-patterned arrays. 185, 1777–1792. e1721.

Chen, Y., Yang, S., Tavormina, J., Tampe, D., Zeisberg, M., Wang, H., Mahadevan, K.K., Wu, C.-J., Sugimoto, H., and Chang, C.-C.J.C.C. (2022b). Oncogenic collagen I homotrimers from cancer cells bind to α3β1 integrin and impact tumor microbiome and immunity to promote pancreatic cancer. 40, 818–834. e819.

Chen, Z., King, W.C., Hwang, A., Gerstein, M., and Zhang, J.J.S.A. (2022c). DeepVelo: Single-cell transcriptomic deep velocity field learning with neural ordinary differential equations. 8, eabq3745.

Chizat, L., Peyré, G., Schmitzer, B., and Vialard, F.-X.J.J.o.F.A. (2018). Unbalanced optimal transport: Dynamic and Kantorovich formulations. 274, 3090–3123.

Cuturi, M.J.A.i.n.i.p.s. (2013). Sinkhorn distances: Lightspeed computation of optimal transport. 26.

Dagogo-Jack, I., and Shaw, A.T.J.N.r.C.o. (2018). Tumour heterogeneity and resistance to cancer therapies. 15, 81–94.

de Visser, K.E., and Joyce, J.A.J.C.C. (2023). The evolving tumor microenvironment: From cancer initiation to metastatic outgrowth. 41, 374–403.

Evans, M.K., Matsui, Y., Xu, B., Willis, C., Loome, J., Milburn, L., Fan, Y., Pagala, V., and Peng, J.C.J.N.c. (2020). Ybx1 fine-tunes PRC2 activities to control embryonic brain development. 11, 4060.

Farrell, J.A., Wang, Y., Riesenfeld, S.J., Shekhar, K., Regev, A., and Schier, A.F.J.S. (2018). Single-cell reconstruction of developmental trajectories during zebrafish embryogenesis. 360, eaar3131.

Ganesh, K., and Massague, J.J.N.m. (2021). Targeting metastatic cancer. 27, 34–44.

Grün, D., Muraro, M.J., Boisset, J.-C., Wiebrands, K., Lyubimova, A., Dharmadhikari, G., van den Born, M., Van Es, J., Jansen, E., and Clevers, H.J.C.s.c. (2016). De novo prediction of stem cell identity using single-cell transcriptome data. 19, 266–277.

Gupta, R., Malvi, P., Parajuli, K.R., Janostiak, R., Bugide, S., Cai, G., Zhu, L.J., Green, M.R., and Wajapeyee, N.J.P.o.t.N.A.o.S. (2020). KLF7 promotes pancreatic cancer growth and metastasis by up-regulating ISG expression and maintaining Golgi complex integrity. 117, 12341–12351.

Haghverdi, L., Buettner, F., and Theis, F.J.J.B. (2015). Diffusion maps for high-dimensional single-cell analysis of differentiation data. 31, 2989–2998.

Hausser, J., and Alon, U.J.N.R.C. (2020). Tumour heterogeneity and the evolutionary trade-offs of cancer. 20, 247–257.

Huang, Y., Lin, L., Shen, Z., Li, Y., Cao, H., Peng, L., Qiu, Y., Cheng, X., Meng, M., and Lu, D.J.A.J.o.C.R. (2020). CEBPG promotes esophageal squamous cell carcinoma progression by enhancing PI3K-AKT signaling. 10, 3328.

Jacomy, M., Venturini, T., Heymann, S., and Bastian, M.J.P.o. (2014). ForceAtlas2, a continuous graph layout algorithm for handy network visualization designed for the Gephi software. 9, e98679.

Ji, Z., and Ji, H.J.N.a.r. (2016). TSCAN: Pseudo-time reconstruction and evaluation in single-cell RNA-seq analysis. 44, e117–e117.

La Manno, G., Soldatov, R., Zeisel, A., Braun, E., Hochgerner, H., Petukhov, V., Lidschreiber, K., Kastriti, M.E., Lönnerberg, P., and Furlan, A.J.N. (2018). RNA velocity of single cells. 560, 494–498.

Li, H., Handsaker, B., Wysoker, A., Fennell, T., Ruan, J., Homer, N., Marth, G., Abecasis, G., and Durbin, R.J.B. (2009). 1000 Genome Project Data Processing Subgroup. 2009. The sequence alignment/map format and samtools. 25, 2078–2079.

Lust, K., Maynard, A., Gomes, T., Fleck, J.S., Camp, J.G., Tanaka, E.M., and Treutlein, B.J.S. (2022). Single-cell analyses of axolotl telencephalon organization, neurogenesis, and regeneration. 377, eabp9262.

McInnes, L., Healy, J., and Melville, J.J.a.p.a. (2018). Umap: Uniform manifold approximation and projection for dimension reduction.

Nitzan, M., Karaiskos, N., Friedman, N., and Rajewsky, N.J.N. (2019). Gene expression cartography. 576, 132–137.

Qin, P., Pang, Y., Hou, W., Fu, R., Zhang, Y., Wang, X., Meng, G., Liu, Q., Zhu, X., and Hong, N.J.C.d. (2021). Integrated decoding hematopoiesis and leukemogenesis using single-cell sequencing and its medical implication. 7, 2.

Qiu, X., Mao, Q., Tang, Y., Wang, L., Chawla, R., Pliner, H.A., and Trapnell, C.J.N.m. (2017). Reversed graph embedding resolves complex single-cell trajectories. 14, 979–982.

Qiu, X., Zhang, Y., Martin-Rufino, J.D., Weng, C., Hosseinzadeh, S., Yang, D., Pogson, A.N., Hein, M.Y., Min, K.H.J., and Wang, L.J.C. (2022). Mapping transcriptomic vector fields of single cells. 185, 690–711. e645.

Ranzoni, A.M., Tangherloni, A., Berest, I., Riva, S.G., Myers, B., Strzelecka, P.M., Xu, J., Panada, E., Mohorianu, I., and Zaugg, J.B.J.C.s.c. (2021). Integrative single-cell RNA-seq and ATAC-seq analysis of human developmental hematopoiesis. 28, 472–487. e477.

Saunders, C.T., Wong, W.S., Swamy, S., Becq, J., Murray, L.J., and Cheetham, R.K.J.B. (2012). Strelka: accurate somatic small-variant calling from sequenced tumor–normal sample pairs. 28, 1811–1817.

Schiebinger, G., Shu, J., Tabaka, M., Cleary, B., Subramanian, V., Solomon, A., Gould, J., Liu, S., Lin, S., and Berube, P.J.C. (2019). Optimal-transport analysis of single-cell gene expression identifies developmental trajectories in reprogramming. 176, 928–943. e922.

Sgubin, M., Pegoraro, S., Pellarin, I., Ros, G., Sgarra, R., Piazza, S., Baldassarre, G., Belletti, B., Manfioletti, G.J.C.D., and Disease (2022). HMGA1 positively regulates the microtubule-destabilizing protein stathmin promoting motility in TNBC cells and decreasing tumour sensitivity to paclitaxel. 13, 429.

Sikder, H.A., Devlin, M.K., Dunlap, S., Ryu, B., and Alani, R.M.J.C.c. (2003). Id proteins in cell growth and tumorigenesis. 3, 525–530.

Street, K., Risso, D., Fletcher, R.B., Das, D., Ngai, J., Yosef, N., Purdom, E., and Dudoit, S.J.B.g. (2018). Slingshot: cell lineage and pseudotime inference for single-cell transcriptomics. 19, 1–16.

Titouan, V., Courty, N., Tavenard, R., and Flamary, R. (2019). Optimal transport for structured data with application on graphs. Paper presented at: International Conference on Machine Learning (PMLR).

Villani, C. (2009). Optimal transport: old and new, Vol 338 (Springer).

Wang, Y., Lin, L., Lai, H., Parada, L.F., and Lei, L.J.D.D. (2013). Transcription factor Sox11 is essential for both embryonic and adult neurogenesis. 242, 638–653.

Wei, X., Fu, S., Li, H., Liu, Y., Wang, S., Feng, W., Yang, Y., Liu, X., Zeng, Y.-Y., and Cheng, M.J.S. (2022). Single-cell Stereo-seq reveals induced progenitor cells involved in axolotl brain regeneration. 377, eabp9444.

Winkler, J., Abisoye-Ogunniyan, A., Metcalf, K.J., and Werb, Z.J.N.c. (2020). Concepts of extracellular matrix remodelling in tumour progression and metastasis. 11, 5120.

Wolf, F.A., Hamey, F.K., Plass, M., Solana, J., Dahlin, J.S., Göttgens, B., Rajewsky, N., Simon, L., and Theis, F.J.J.G.b. (2019). PAGA: graph abstraction reconciles clustering with trajectory inference through a topology preserving map of single cells. 20, 1–9.

Wu, L., Yan, J., Bai, Y., Chen, F., Zou, X., Xu, J., Huang, A., Hou, L., Zhong, Y., and Jing, Z.J.C.R. (2023). An invasive zone in human liver cancer identified by Stereo-seq promotes hepatocyte–tumor cell crosstalk, local immunosuppression and tumor progression. 1–19.

Zappia, L., Phipson, B., and Oshlack, A.J.G.b. (2017). Splatter: simulation of single-cell RNA sequencing data. 18, 174.

Zhang, T., Liu, D., Wang, Y., Sun, M., and Xia, L.J.F.i.O. (2021). The E-Twenty-Six Family in Hepatocellular Carcinoma: Moving into the Spotlight. 10, 620352.

