## Supplementary material for "Inferring cell trajectories of spatial transcriptomics via optimal transport analysis": Figure S1

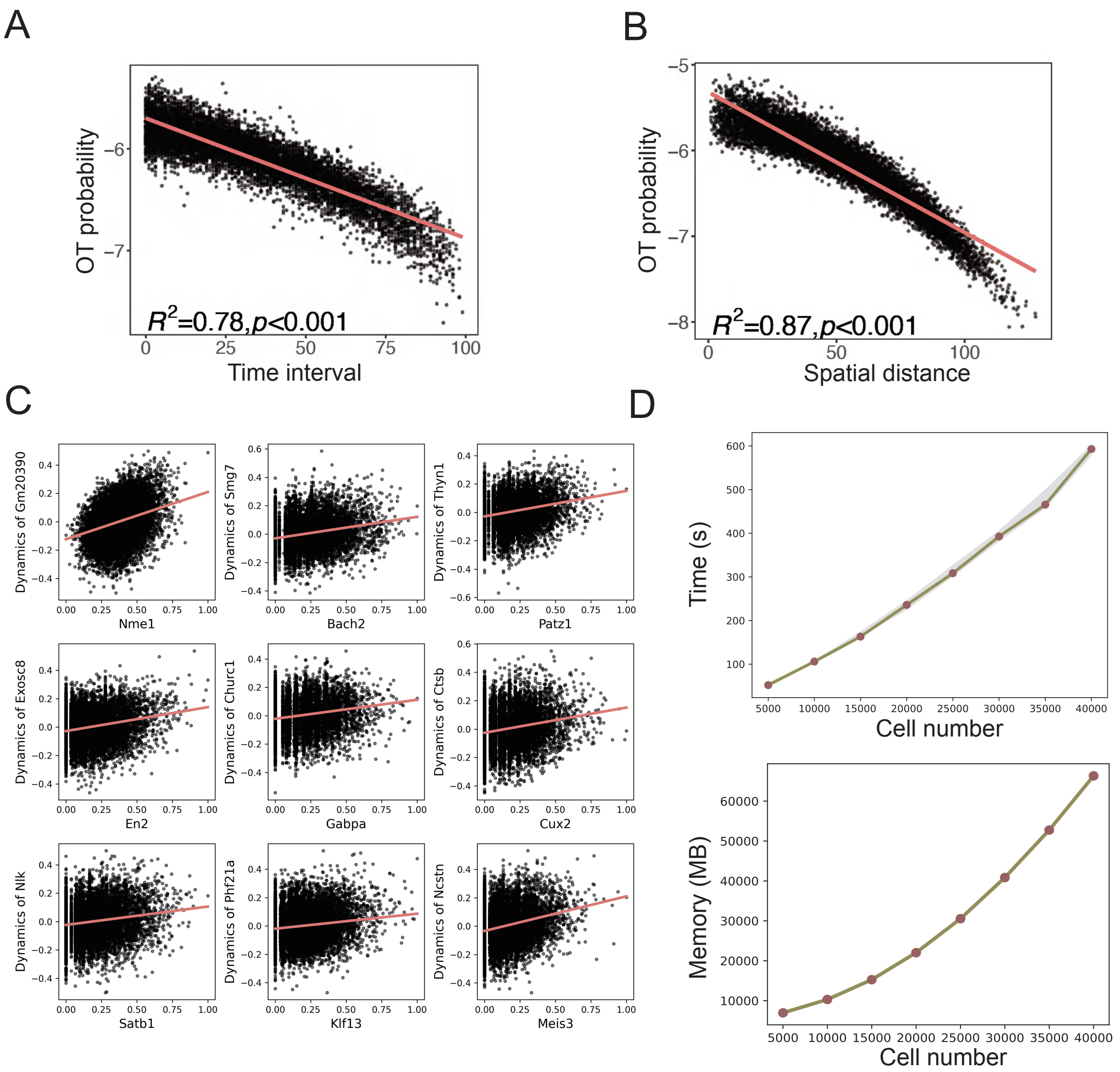


**Figure S1 spaTrack effectively and efficiently captures the cell transitions and driven factors.**

A. Connection between the transition probability and time intervals. The SC data with time intervals (1-100) was simulated using Splatter. The plots were combined from 100 repeats of simulation and computation. Parameters of simulation are assigned as nGenes=3000, batchCells=3000, mean.shape=0.6, mean.rate=0.3, bcv.common=0.2, dropout.mid=0, dropout.shape=-1, out.prob=0.05, de.prob=1, method = "paths", path.from=c(0), path.length=c(100), path.skew=c(0.5), path.nonlinearProb=0.1.

B. Connection between the transition probability and spatial distance. ST data was simulated following two steps: 1) simulated SC data of one group using Splatter; 2) assigned the 3000 cells of each simulation to a 5000 μm × 5000 μm square, assuming each cell taking up a 50 μm × 50 μm spot. The plots were combined from 100 repeats of simulation and computation.

C. Regulatory connection between TFs and targets inferred by spaTrack. A neural network framework is implanted in the algorithm, with expression profile of genes at $t_1$ time as input layer, prediction of TF expression of $t_0$ time as output layer. TF-target pairs with high weights were plotted to show their regulatory connection.

D. Time and memory cost of spaTrack in trajectory inference. We applied spaTrack on different cell population size ranging from 5k to 40k. Under a standard CPU thread (Intel(R) Xeon(R) CPU E5-2650 v4 @ 2.20GHz), spaTrack required only minutes to finish the computation of 5k – 400k cells (with 20,000 features). The memory load depends seriously on the population size. With 20,000 features, it followed an exponential growth with 6.9 GB for 5k cells.

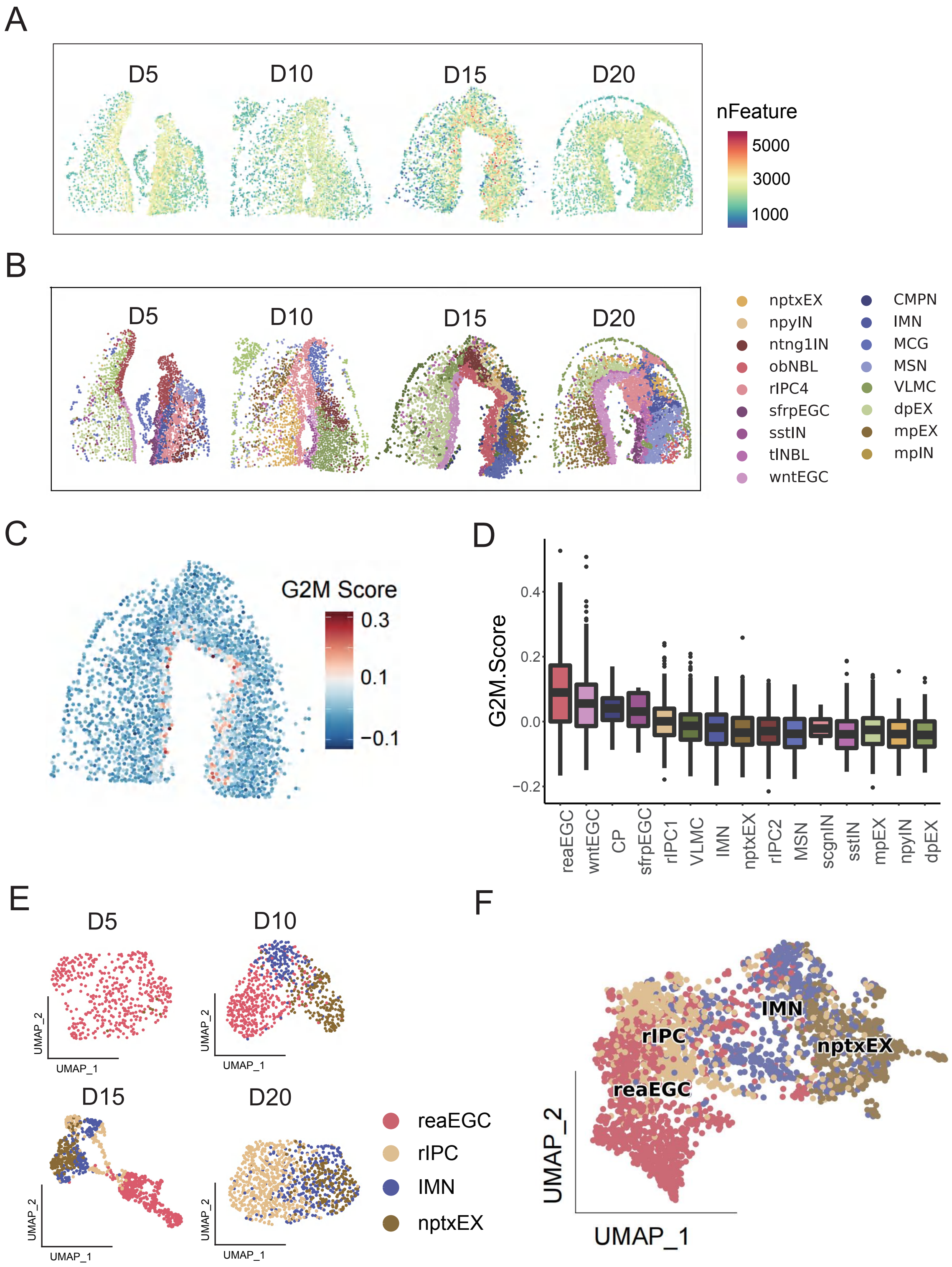


**Figure S2 Fine local trajectories of axolotl telencephalon regeneration**

A. Spatial distribution of feature number of the ST data, sampled from the regenerative stages of axolotl telencephalon at 5 days (D5), 10 days (D10), 15 days (D15), and 20 days (D20) after injury.

B. Spatial distribution of cell types of each stage.

C. Spatial distribution of G2M score in ST data of D15.

D. G2M score of each cell type of D15.

E. Sparse cell types and cell continuity of regenerative cells in single sample.

F. Integration of regenerative cells from multiple ST samples.

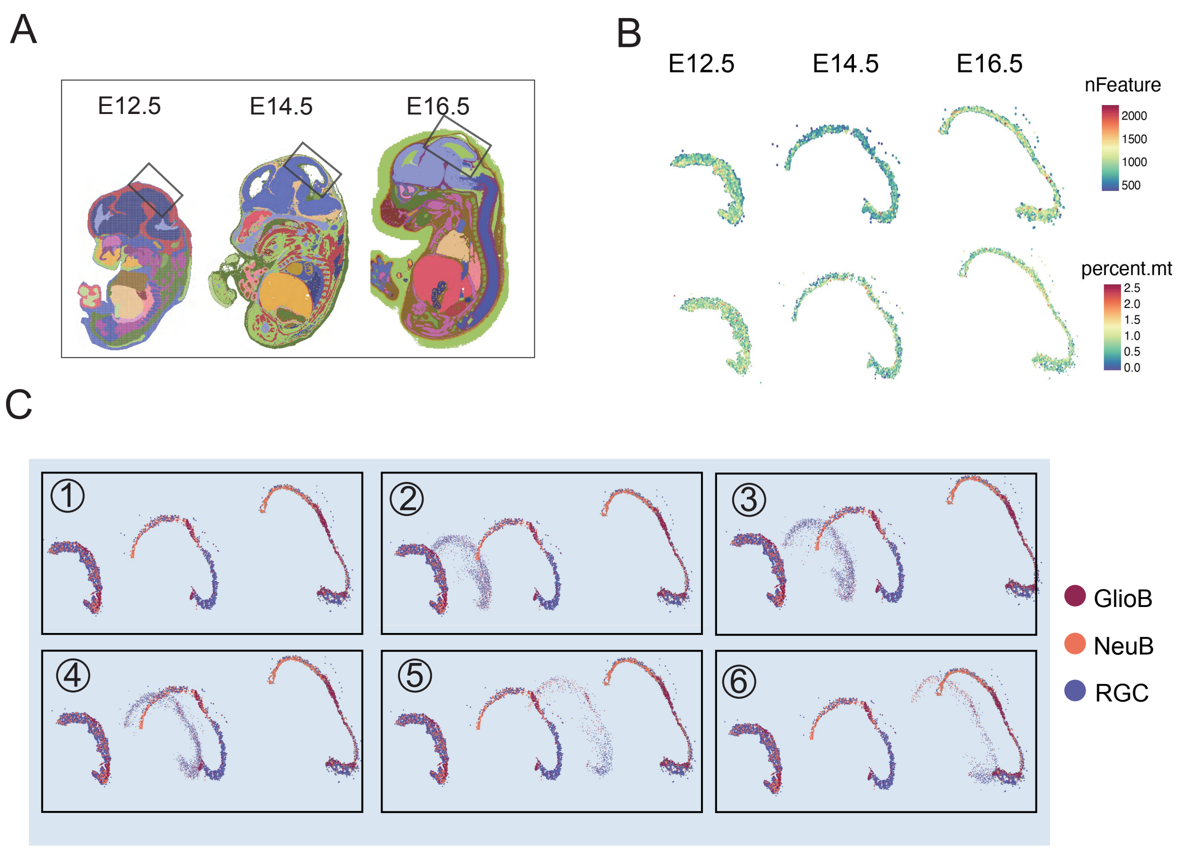


**Figure S3 Tracing neuron cells across mouse embryos of a time series**

A. Collections of mouse embryo ST data and their dorsal midbrain regions, at day 12.5 (E12.5), 14.5 (E14.5), and 16.5 (E16.5) of embryonic stages.

B. Spatial distribution of feature number and proportion of mitochondrial reads in the ST data of dorsal midbrain.

C. Screen shots of the dynamic visualization of the RGC differentiation across different sections. The animation display could be found at https://spatrack.readthedocs.io/en/latest/index.html

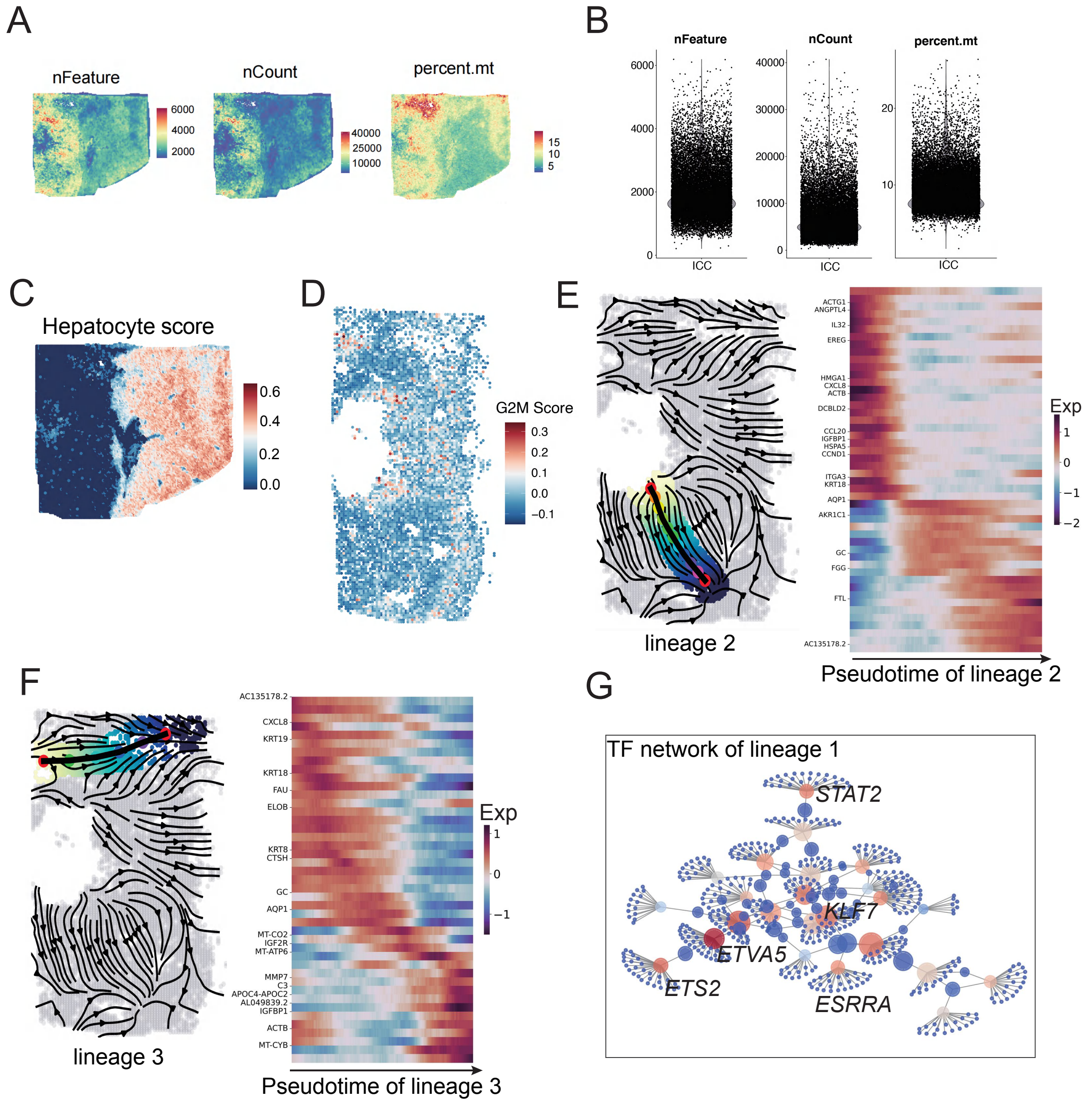


**Figure S4 Recovering the spatial trajectories of tumor expansion**

A. Spatial distribution of feature number, UMI number, and mitochondrial reads of the ST data of a primary tumor of intrahepatic cholangiocarcinoma (ICC).

B. Violin plots of feature number, UMI number, and mitochondrial reads of the ST data.

C. Spatial distribution of hepatocytes of the ST data.

D. Spatial distribution of G2M score of the tumor cells.

E. Optimal path of the lineage 2 (P0-P3-P4) and pseudotime-dependent genes. These genes were screened by fitting a generalized additive model. Pseudotime is obtained by projecting a cell to the optimal path and the arc length of projected position is normalized as a relative pseudotime.

F. Optimal path of the lineage 3 (P0-P5-P6) and pseudotime-dependent genes.

G. Regulatory network underlying tumor expansion of lineage 1.

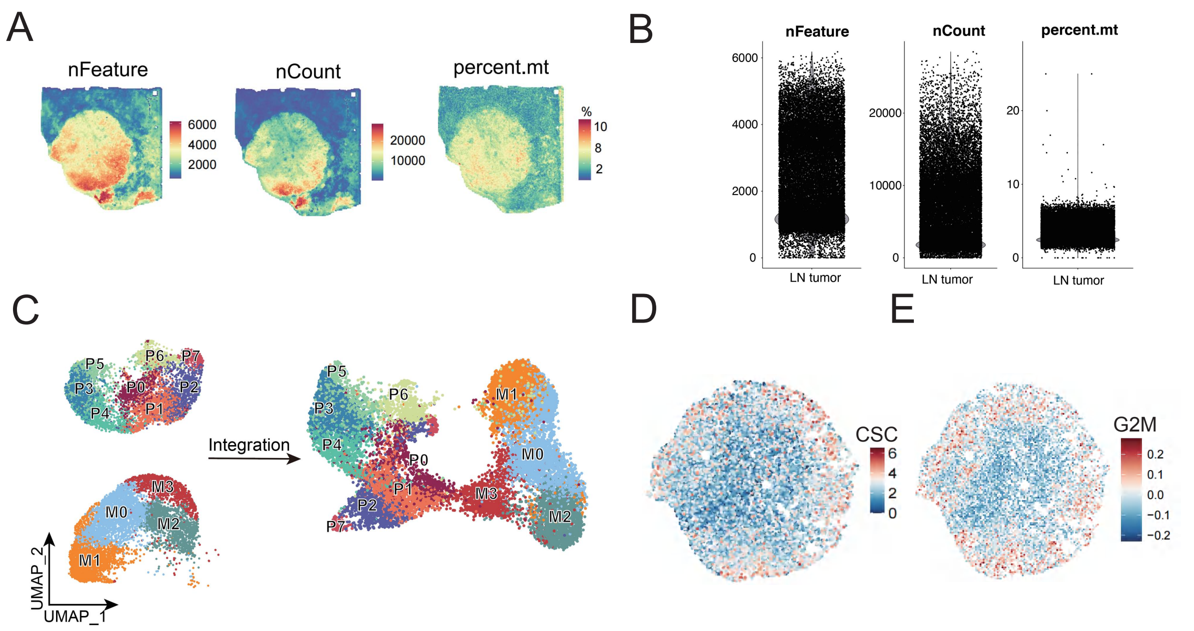


**Figure S5 Tracing tumor metastasis**

A. Spatial distribution of feature number, UMI number, and mitochondrial reads of the ST data of the metastatic tumor in lymph node, corresponding with the primary ICC in Figure 5A.

B. Violin plots of feature number, UMI number, and mitochondrial reads of the ST data of the metastatic tumor.

C. Integration of primary tumor and its metastatic tumor with Harmony in a SC manner.

D. Spatial distribution of the expression of cancer stem cell markers.

E. Spatial distribution of G2M score.

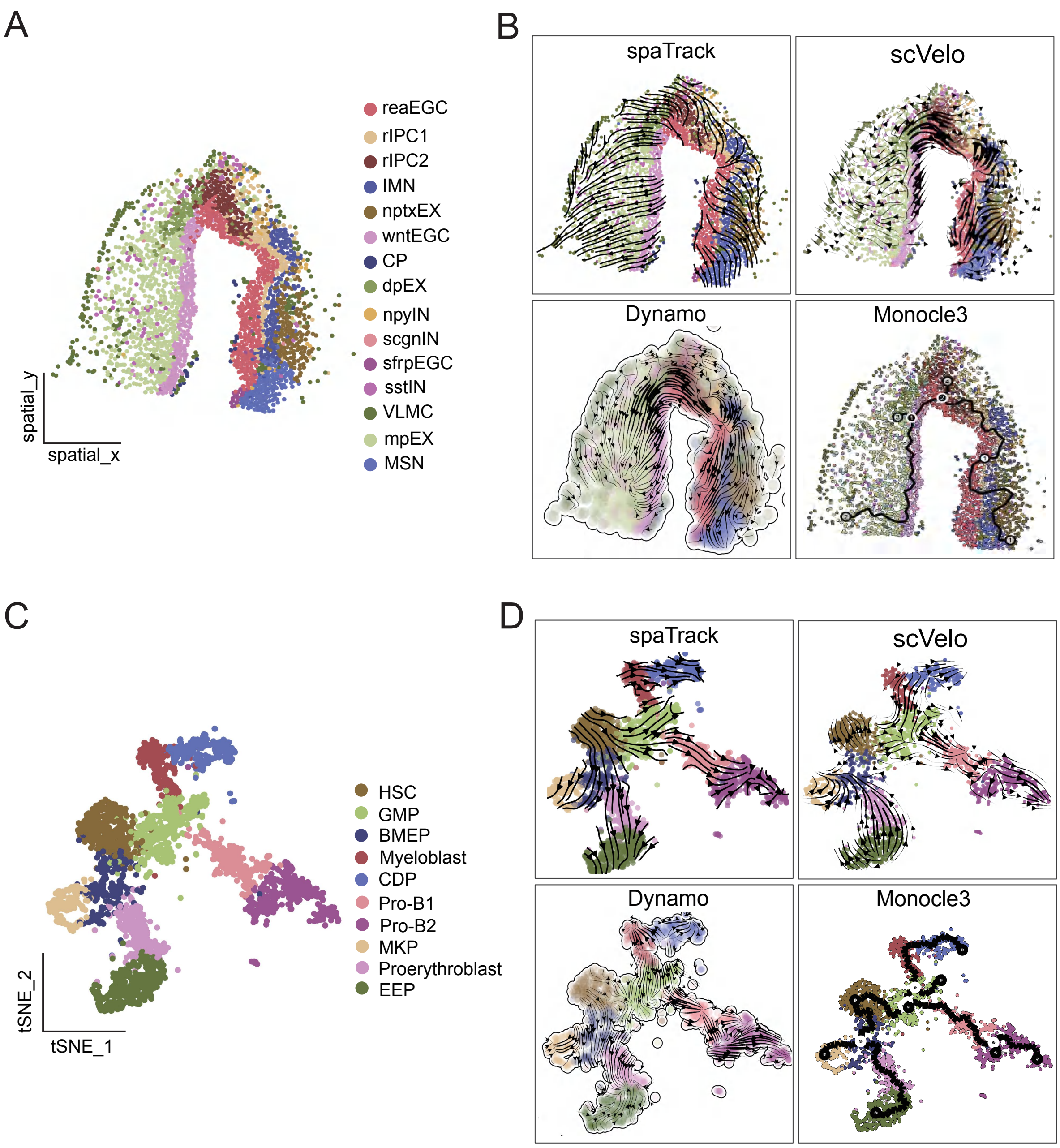


**Figure S6 Comparison of performance between spaTrack and other methods in empirical data with complex differentiation topology.**

A. A ST data with annotated cell types was used for the comparison of TI performance between methods. The sample of axolotl telencephalon was collected at 15th day after injury. The regeneration is spatially organized as three lineages as described in our Results section.

B. TI of spaTrack, scVelo, Dynamo, and Monocle3 on the ST data of axolotl telencephalon regeneration.

C. A SC data of primary human hematopoietic stem and progenitor cells (HSPCs) with annotated cell types was used for the comparison of TI performance between methods.

Human hematopoiesis is well studied as a hierarchical and multiple-branched process. The complex topology and various progenitors challenge the common TI methods.

D. Comparing the TI of spaTrack, scVelo, Dynamo, and Monocle3 on the SC data of HSPCs.
